# β-hydroxybutyrate improves convulsions in mice carrying the *GRIN1* Y647S/+ pathogenic variant

**DOI:** 10.1101/2024.08.21.608984

**Authors:** Megan T. Sullivan, Patrick Tidball, Yuanye Yan, Katheron Intson, Wenjuan Chen, Yuchen Xu, Sridevi Venkatesan, Wendy Horsfall, John Georgiou, Peter S.B. Finnie, Robert E. McCullumsmith, Evelyn K. Lambe, Stephen F. Traynelis, Ali Salahpour, Hongjie Yuan, Graham L. Collingridge, Amy J. Ramsey

## Abstract

*GRIN1*-related neurodevelopmental disorder (*GRIN1*-NDD) is characterized by clinically significant variation in the *GRIN1* gene, which encodes the obligatory GluN1 subunit of N-methyl-D-aspartate receptors (NMDARs). The identified p.Tyr647Ser (Y647S) variant is carried by a 34-year-old female with seizures and intellectual disability. This study builds upon initial *in vitro* investigations of the functional impacts of this variant in the SYTANLAAF domain of the GluN1 M3 helix and examines its *in vivo* consequences in a mouse model.

To investigate *in vitro* functional impacts of NMDARs containing GluN1-Y647S variant subunits, GluN1-Y647S was co-expressed with wildtype GluN2A or GluN2B subunits in *Xenopus laevis* oocytes and HEK cells. *Grin1*^Y647S/+^ mice were created by CRISPR-Cas9 endonuclease-mediated transgenesis and the molecular, electrophysiological, and behavioural consequences of the variant were examined. Additionally, de-identified patient data were collected to examine the representative nature of *Grin1*^Y647S/+^ mice in modelling specific aspects of patient symptomology.

*In vitro*, NMDARs containing GluN1-Y647S showed altered sensitivity to endogenous agonists and negative allosteric modulators, and reduced cell surface trafficking. *Ex vivo*, *Grin1*^Y647S/+^ mice displayed a reduction in whole brain GluN1 levels and a deficiency in NMDAR-mediated synaptic transmission in the hippocampus. Behaviourally, *Grin1*^Y647S/+^ mice exhibited altered vocalizations, muscle strength, sociability, and problem-solving, as well as spontaneous convulsions that were ameliorated with supplementation of β-hydroxybutyrate (BHB), an endogenously produced ketone body.

The Y647S variant confers a complex *in vivo* phenotype, which reflects largely diminished properties of NMDAR function. As a result, *Grin1*^Y647S/+^ mice display atypical behaviour in domains relevant to the clinical characteristics of *GRIN1*-NDD and the individual carrying the variant, which allowed for the identification of BHB supplementation as a potential anti-convulsant treatment. Ultimately, the characterization of *Grin1*^Y647S/+^ mice accomplished in the present work, expands our understanding of the mechanisms underlying *GRIN1*-NDD and provides a foundation for the continued development of novel therapeutics.

## Introduction

*GRIN-*related neurodevelopmental disorder (*GRIN-*NDD) is characterized by clinically significant variation in genes coding for the subunits of the N-methyl-D-aspartate receptor (NMDAR).^1–3^ The *GRIN1* gene encodes the obligatory GluN1 subunit^4^ which is required for proper function of all NMDARs.^5–7^ NMDARs flux calcium when activated by concurrent occupation of glycine and glutamate sites on the GluN1 and GluN2 subunits, respectively, together with membrane depolarization.^6,8,9^ These receptors are crucial to cognitive processes like learning and memory through mediation of brain plasticity.^10–12^

*GRIN1* variants can be familial in origin but are typically *de novo* missense variants.^3^ With fewer than 100 published cases, the prevalence of *GRIN1*-related neurodevelopmental disorder (*GRIN1*-NDD) in the general population has yet to be established,^3^ but is estimated to be ∼5 per 100,000 births.^13^ These patients have a wide range of symptoms of varying severity. Nearly all present with some form of developmental delay, atypical cognition, and intellectual disability, while two thirds of patients present with epilepsy, stereotyped movements, and abnormal eye movements resembling oculogyric crisis.^14^ Other observed symptoms include muscular hypotonia, self-injurious behaviours, and autism spectrum disorder (ASD).^3^ Current treatment of *GRIN1*-NDD involves symptomatic management of these disease manifestations.^3^ For patients with epilepsy, this includes conventional anti-seizure medications, of which many *GRIN1*-NDD patients are refractory to treatment^3^ and in some cases use of the ketogenic diet.^15^ This consists of a high-fat, low protein and carbohydrate diet which leads to the production of ketone bodies, principally β-hydroxybutyrate (BHB), which is hypothesized to mediate the diet’s anticonvulsant effects.^16^

The diversity of phenotypes observed across *GRIN1*-NDD patients has been attributed, in part, to the multitude of unique *GRIN1* gene variants. While variants are often unique to each patient, they tend to cluster near or within functionally important receptor domains, including the agonist-binding domains and transmembrane domain that forms the intrinsic ion channel pore of the NMDAR.^1,2,14,17–19^ Individual variants have been found to enhance or diminish various properties of NMDARs and can be broadly classified as gain-of-function (GoF) or loss-of-function (LoF).^20^ While the location of certain gene variants has been linked to specific phenotypic features among *GRIN1*-NDD patients, including cortical malformation,^3,17^ such relationships remain ill-defined. Clinical differences are even inconsistent between patients possessing *GRIN1* variants categorized as NMDAR “GoF” or “LoF”, posing a challenge to the development and delivery of *GRIN1*-NDD therapeutics. Therefore, in-depth study of individual variants is necessary to determine the unique functional consequences of variation in the NMDAR, to establish predictive variant-function relationships, and to ultimately guide patient treatment.^21^

In this study we examine the p.Tyr647Ser (Y647S) variant which is located in the transmembrane domain M3 helix of GluN1 and has been identified in two patients, one of which is a 34-year-old female with seizures and intellectual disability.^2,22^ This variant falls within the highly conserved 646-654 residues of M3 known as the Lurcher (SYTANLAAF) motif. The motif is thought to be crucial to the gating function of the receptor^14,23^ and potentially to the trafficking of NMDARs to the cell surface.^24^ *In vitro*, the Y647S variant reduces maximal glutamate-inducible current amplitudes in heterologous expression systems, and accompanying *in silico* models predict that the Y647S variant destabilizes quaternary structure of the transmembrane domain, disrupting NMDAR assembly.^2^ The Y647S variant also reduces the surface delivery of NMDARs when expressed in HEK293 cells or in rat hippocampal neurons.^25^ Together, these studies point to the Y647S variant as generating an NMDAR LoF phenotype.^2,20^ Nevertheless, Xu *et al.*^26^ classified the Y647S variant as one of indeterminant status,^20^ as NMDARs containing GluN1-Y647S display diminished open probability and surface expression (as noted in previous studies)^25^ alongside increases in glutamate and glycine potency, compared to NMDARs containing wildtype GluN1.

Here, we build upon the initial *in vitro* investigations of the Y647S patient variant by examining the molecular, electrophysiological, and behavioural consequences of the variant using a novel mouse model. Additionally, we further describe the clinical features of the 34-year-old female patient carrying the Y647S variant to validate the *Grin1*^Y647S/+^ mouse as a representative model of key features of patient symptomology. Our comprehensive characterization reveals the Y647S variant confers a complex *in vivo* phenotype, which reflects largely diminished properties of NMDAR function. *Grin1^Y^*^647S/+^ mice displayed neuroanatomical and behavioural abnormalities reminiscent of the individual carrying this variant and the broader clinical characteristics of *GRIN1*-NDD. Notably, *Grin1*^Y647S/+^ mice exhibited spontaneous convulsions and the present work identified BHB supplementation as a potential anti-convulsant treatment. Our findings underscore the utility of this model in future studies seeking to further understand the pathological mechanisms underlying NMDAR *GRIN* variation and to test additional novel therapies.

## Materials and methods

### Collection of de-identified patient data

Consent was obtained to share de-identified patient data with collaborating researchers according to the Declaration of Helsinki and under a protocol approved by the University of Toronto Research Ethics Board (#00047662; Use of patient medical information in the study of *GRIN1*-Related Neurodevelopmental Disorder). Upon provision of written informed consent, the caregiver of the individual carrying the Y647S variant was provided with the study questionnaire. The questionnaire consisted of five sections: general information, diagnosis, current symptomology & treatment regimens, previously tried medications and personal information. Questions inquired about patient age, sex, clinical presentation at birth, the process of receiving a *GRIN1*-NDD diagnosis, symptomology, current treatment regimens and responses to previously tried medications, as well as patient’s strengths, challenges, and unmet treatment needs.

### Molecular biology and two-electrode voltage clamp current recordings

The complementary DNA (cDNA) encoding the human GluN1-1a (referred to as GluN1; NM_007327.3) was utilized to generate the human *GRIN1*_p.Y647S variant by site-directed mutagenesis using the QuikChange protocol (Stratagene La Jolla, CA, USA) in the plasmid vector pCI neo as described previously.^27^ Wild type (WT) or variant GluN1 subunit was co-expressed with human GluN2A (NM_000833.4) or human GluN2B (NM_000834.4) in pCI neo. The cDNA was linearized using DpnI (ThermoFisher, Waltham, MA) restriction digestion at 37°C for 1 hour. Complementary RNA (cRNA) was synthesized *in vitro* using the mMessage mMachine T7 kit (Ambion, Austin, TX, USA) from linearized WT and variant cDNA (DpnI restriction digestion at 37°C for 1 hour; ThermoFisher, Waltham, MA).

Unfertilized *Xenopus laevis* stage VI oocytes were prepared from commercially available ovaries (Xenopus one Inc, Dexter, MI, USA) as described previously.^28^ *Xenopus laevis* oocytes were injected with a 1:2 ratio of GluN1:GluN2A or GluN1:GluN2B cRNA and maintained in normal Barth’s solution (in mM; 88 NaCl, 2.4 NaHCO_3_, 1 KCl, 0.33 Ca(NO_3_)_2_, 0.41 CaCl_2_, 0.82 MgSO_4_, 10 HEPES, pH 7.4 with NaOH; supplemented with 100 μg/mL gentamycin, 40 μg/mL streptomycin) at 19°C for 2-3 days before recordings.

Two-electrode voltage clamp (TEVC) current recordings were performed at room temperature (23°C) with the extracellular recording solution containing (in mM) 90 NaCl, 1 KCl, 10 HEPES, 0.5 BaCl_2_, and 0.01 EDTA (pH 7.4 with NaOH) as previously described.^27,29^ The voltage electrode and current electrode were filled with 0.3 M KCl and 3 M KCl, respectively. Current responses from oocytes were recorded under voltage clamp at holding potential (V_HOLD_) -40 mV except for experiments that involved changes in the concentration of extracellular Mg^2+^ and Zn^2+^, for which case V_HOLD_ was set at -60 mV and -20 mV, respectively. Glutamate concentration-response curves were recorded in the presence of saturating glycine, and glycine-concentration response curves were recorded in the presence of saturating glutamate. The concentration-response relationships for glutamate and glycine were fitted by *Equation 1*,

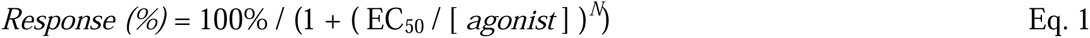

The concentration-response relationship for inhibition by Mg^2+^, Zn^2+,^ and H^+^ was recorded in the presence of saturating concentrations of both glutamate and glycine. Zn^2+^ concentration was buffered as previously described.^20,30^ Concentration-response relationship for Mg^2+^, Zn^2+^ and extracellular pH (proton concentration) were fitted by *Equation 2*,

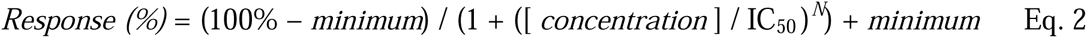

where EC_50_ is the concentration of the agonist (glutamate or glycine) that generates a half-maximal effect, IC_50_ is the concentration of the inhibitor/blocker that causes a half-maximal effect, *minimum* is the degree of residual response at a saturating concentration of the inhibitor/blocker, *N* is the Hill slope, and *Response* is indicated as a percentage of the fitted maximum.

The channel open probability (*P_OPEN_*) was calculated from the degree *Potentiation* measured in MTSEA (2-aminoethyl methanethiol sulfonate hydrobromide, Toronto Research Chemicals, Ontario, Canada) using *Equation 3* ^31^:

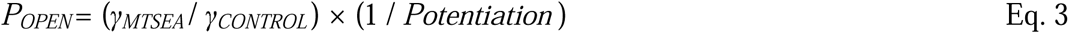

where γ*_MTSEA_* and γ*_CONTROL_* were the single channel chord conductance values estimated from GluN1/GluN2A receptors and fold *Potentiation* was defined as the ratio of current in the presence of MTSEA to current in the absence of MTSEA, and γ*_MTSEA_* / γ*_CONTROL_* was 0.67.^31^ MTSEA solution was made fresh from powder stock and used within 30 minutes.

### Whole cell voltage clamp recordings and beta-lactamase assay

HEK293 cells (CRL 1573, ATCC, Manassas, VA, USA) were maintained in standard DMEM/Gluta-Max media (Thermo Fisher, Waltham, MA, USA) with 10% dialyzed fetal bovine serum (R&D Systems, Minneapolis, MN, USA) supplemented with 10 U/mL penicillin and 10 µg/mL streptomycin (Thermo Fisher, Waltham, MA, USA) at 37°C and 5% CO_2_. The calcium phosphate method was used to transiently transfect the cells with plasmid cDNA encoding green fluorescent protein (GFP) and NMDAR GluN subunits (5:1:1 for GFP:GluN1:GluN2A or variant GluN1-Y647S, 1:1:1 for GFP:GluN1:GluN2B or variant GluN1-Y647S), as previously described.^32^ Whole cell voltage clamp (WCVC) recordings^27,32^ were performed on the transfected HEK293 cells 24-48 hours post-transfection with external recording solution that contained (in mM) 150 NaCl, 10 HEPES, 30 D-mannitol, 3 KCl, 1 CaCl_2_, and 0.01 EDTA (pH 7.4). The patch electrodes (resistance 3-5 MΩ) were prepared from thin-walled glass micropipettes (TW150F-4, World Precision Instruments, Sarasota, FL, USA) and filled with internal solution (in mM; 110 D-gluconate, 110 CsOH, 30 CsCl, 5 HEPES, 4 NaCl, 0.5 CaCl_2_, 2 MgCl_2_, 5 BAPTA, 2 NaATP and 0.3 NaGTP, pH 7.35). The current response was recorded (V_HOLD_: -60 mV) using an Axopatch 200B amplifier (Molecular Devices, Union City, CA, USA) at room temperature (23°C).

For the β-lactamase surface expression assay, HEK293 cells were plated in 96-well plates (50,000 cells per well) and transiently transfected using Fugene6 (Promega, Madison, WI) with cDNA encoding a fusion protein of β-lac-WT GluN1 or β-lac-GluN1-Y647S and wild type GluN2B.^33^ Eight wells were transfected with surface and total expression activities measured in 4 wells each 24 h post-transfection. The absorbance at 486 nm was measured by a microplate reader at 30 every min for 30 min. The ratio of surface (unlysed cells) to total (lysed cells) β-lactamase expression was measured for the blac-GluN1-Y647S and compared to blac-GluN1-WT as described previously.^33^ All chemicals were purchased from Sigma-Aldrich unless otherwise stated.

### Mice and genotyping

Mice were housed with littermates on a 12-hour light-dark cycle with food and water available *ad libitum*. All experimentation was conducted during the light phase. Animal housing and experiments were conducted in accordance with the Canadian Council on Animal Care, the University of Toronto Faculty of Medicine and Pharmacy Animal Care Committee and/or The Centre for Phenogenomics Animal Care Committee.

*Grin1*^Y647S/+^ (Y647S/+) mice were created via CRISPR-Cas9 endonuclease-mediated transgenesis. Founder mice were identified by Sanger sequencing and bred to a C57BL/6J background to generate Wildtype (WT, +/+) and Y647S/+ mice. Mice were genotyped using two separate PCRs for each allele: the WT and Y647S variant allele (which is a two base pair difference). Therefore, each PCR reaction is completed with either the WT primers or Y647S variant allele amplification primers (339-bp amplicon) (along with a control reaction for PCR); WT - forward: GAT CAT CGT GGC TTC CTA reverse: CTC AGC TGC ACT CTC ATA AT; Y647S - forward: GAT CAT CGT GGC TTC TTC reverse: CTC AGC TGC ACT CTC ATA AT. Touchdown PCR was used with DreamTaq DNA Polymerase (Thermo Fisher Scientific): 2 min at 96 degrees, then each cycle after is 94 degrees for 20 s, annealing for 30 s. The initial 12 cycles are touchdown from 64-58 degrees (0.5 degrees/cycle), with the remaining 28 cycles with an annealing temperature of 58 degrees (40 cycles total), with extension of 30 s for each cycle.

### RNA isolation and RT-qPCR

Mice were euthanized by cervical dislocation and RNA from forebrain was extracted using TRIzol Reagent (Thermo Fisher Scientific). cDNA was synthesized using an Invitrogen Superscript III Reverse Transcriptase kit (Thermo Fisher Scientific) using 1 μg of sample. Quantitative RT-qPCR was performed on a QuantStudio 5 (Applied Biosystems). Melt curve analysis was used to check reaction specificity. All data was normalized to PGK1-2. Primers were as follows: *Grin1*-forward: ACA CCA ACA TCT GGA AGA CAG reverse: CAG TCA CTC CAT CTG CAT ACT T, *Grin2a –* forward: ACC ATT GGG AGC GGG TAC AT reverse: CCT GCC ATG TTG TCG ATG TC, *Grin2b* – forward: TCC GCC GTG AGT CTT CTG reverse: CTG GGT GGT AAA GGG TGG, *PGK1-2* – forward: GGC CTT TCG ACC TCA CGG reverse: GTC CAC CCT CAT CAC GAC.

### Western blot protein analysis

Mice were euthanized by cervical dislocation and total brain protein lysates were prepared by homogenizing right brain hemispheres in 0.32 M HEPES-buffered sucrose and a protease inhibitor cocktail (PMSF, aprotinin, leupeptin, pepstatin and benzamidine). Synaptic plasma membrane fractions were obtained using a discontinuous sucrose gradient as described previously.^34^ Protein concentration of samples was determined using Bradford protein assay using bovine serum albumin standards. Protein samples (10 μg/lane) were separated on a 10% Tris-glycine gel (Thermo Fisher Scientific) and transferred to a PVDF membrane (Millipore). Blocking of non-specific binding was completed in 3% milk in Tween-Tris buffered saline (TBST) for 1 hour. Primary antibodies were diluted in 3% milk in TBST and incubated overnight at 4 degrees. Antibodies used: NMDAR1 (1:1000; Millipore, Cat# AB9864R), NMDAR2A (1:1000, Millipore, Cat# AB1555P), NMDAR2B (1:1000; Millipore, Cat# AB1557P). Secondary goat anti-rabbit antibody (LI-COR Biosciences; Cat# 926-32213) was diluted at 1:5000 in 3% milk in TBST. Blots were normalized to Revert 700 Total protein stain, visualized using the Odyssey M Imager, and relative band intensities were quantified using Empiria Studio Software (LI-COR Biosciences).

### Hippocampal slice electrophysiology

Hippocampal slices were prepared from adult (12–15 week) male and female Y647S/+ mice and their wildtype (+/+) littermate counterparts. Mice were euthanized by cervical dislocation under isoflurane anaesthesia and brains were rapidly extracted and submerged in ice-cold cutting solution composed of (in mM): 205 Sucrose, 26 NaHCO_3_, 10 Glucose, 2.5 KCl, 1.25 NaH_2_PO_4_, 0.5 CaCl_2_, and 5 MgSO_4_ (saturated with 95% O_2_ and 5% CO_2_; pH 7.4). After being allowed to cool for 2 – 3 min, brains were hemisected along the longitudinal fissure and slices of dorsal hippocampus (400 µm) were obtained by sectioning each hemisphere along its sagittal plane using a VT1200S vibratome (Leica). The CA3 region of slices was removed by scalpel cut immediately after slicing. Slices were then allowed to recover for a minimum of 1 h at room temperature in standard artificial cerebrospinal fluid (ACSF) composed of (in mM): 124 NaCl, 26 NaHCO_3_, 10 Glucose, 3 KCl, 1.4 NaH_2_PO_4_, 2 CaCl_2_, and 1 MgSO_4_ (saturated with 95% O_2_ and 5% CO_2_; pH 7.4).

Following recovery, slices were transferred to a submerged-type chamber where they were continuously perfused with ACSF (2.5 mL/min) and maintained at 30 °C for extracellular field potential recordings which were performed blind to genotype. Glass microelectrodes (∼1.5 MΩ) filled with ACSF were used to record fibre volleys (FVs) and field excitatory postsynaptic potentials (fEPSPs) from CA1 stratum radiatum in response to stimulation of the Schaffer collateral/commissural (SC) pathway. Constant current stimulus pulses (100 µs) were generated by an STG4002 stimulus isolator (Multi Channel Systems) and delivered via a platinum/iridium bipolar stimulating electrode (FHC) positioned in the stratum radiatum near the CA1/CA2 border. Signals were amplified using a Multiclamp 700B (Molecular Devices), filtered at 2 kHz, digitized at 40 kHz, and recorded to a personal computer for offline analysis using WinLTP software.^35^

For each slice, stimulus intensity was determined as a function of the threshold required to elicit a visually detectable response. Input/output (I/O) curves were generated by delivering stimuli at multiples of this threshold value (1– 5×). Baseline stimulus intensity for paired-pulse facilitation (PPF) and long-term potentiation (LTP) experiments was set to 3× the threshold value to evoke sub-maximal AMPAR-mediated fEPSPs (AMPAR-fEPSPs). Test pulses were delivered every 30 s, and four consecutive responses were averaged for analysis. PPF was assessed across a range of inter-pulse intervals (50, 100, 150, 200 and 250 ms). NMDAR-mediated fEPSPs (NMDAR-fEPSPs) were recorded in low (0.1 mM) magnesium ACSF in the presence of the AMPAR blocker NBQX (10 µM). For plasticity experiments, LTP was induced after 30 min of stable baseline recording by delivering a ‘compressed’ theta-burst stimulation (TBS) protocol comprising 3 episodes of 5 bursts of 5 pulses at 100 Hz repeated at 5 Hz, with a 10 s interval between each episode.^36^ FVs were quantified by their peak amplitudes and fEPSPs were quantified by their initial slopes. I/O curves from individual slices were quantified by plotting the FV amplitude as a function of stimulus intensity or the fEPSP slope as a function of FV amplitude and taking the slope value of the fitted linear regression line (I/O slope).

### Histology

Adult male and female mice (13-21 weeks) were euthanized by cervical dislocation and brains were fixed in 10% neutral buffered formalin (Fisher Scientific) for 48 hours. Brains were subsequently dehydrated in 70% ethanol, processed, and embedded into paraffin. Sections were cut sagittally 1.525 mm lateral of Bregma at 5 µm thickness and stained with hematoxylin and eosin or Nissl stain. Brain sections were imaged using the Axioscan 7 Slide Scanner (Zeiss) and subsequently analyzed using QuPath v0.3.2^37^ blind to genotype. The number of cortical neurons was counted in a 1.44 mm^2^ region and thickness measurements were completed using line annotations of the neocortex and hippocampus (including individual measurements of CA1, CA2 and dentate gyrus regions), in a manner similar to Amador *et al*.^38^ Contour tracing for cell body area measurements of the hippocampus proper and dentate gyrus was applied using the QuPath wand tool and fine-tuned using the brush tool in a manner similar to Edwards *et al.*^39^

### Early developmental behavioural phenotyping

#### Ultrasonic Vocalizations (USVs)

USVs were collected at postnatal day (P) 6 in mice of both sexes. Pups were taken in random order from their home cages (containing littermates and mother), and each placed individually into a clean empty cage within a transparent sound attenuating chamber (33.5 x 21.5 x 30.5 cm). The three-minute trial began immediately after the pup was placed into the chamber and lid had been closed. USV recordings were captured using an Avisoft UltraSoundGate condenser microphone CM16 (Avisoft Bioacoustics, Berlin, Germany), sensitive to 10-180 kHz. The microphone was mounted on a clamp located within the sound attenuating chamber approximately 19 cm away from the pup being recorded. Vocalizations were recorded using Avisoft Recorder Software at 250 kHz sampling rate and 16-bit resolution. Analysis was completed using DeepSqueak software v3.1.0^40^ by an experimenter blinded to genotype. This was done using the Mouse Detector YOLO R2 Neural network with 30 kHz lower frequency and 110 kHz upper frequency cutoffs,^41,42^ with a total analysis length of 180 seconds. Score threshold was set to 0, meaning all detections were saved to the file. Calls were visualized using the “Load call” function, and a manual selection review was conducted by an experienced experimenter.

Following collection of USVs, the righting reflex was tested by placing the pup on its back in a clean cage and recording the time it takes for the pup to return to the prone position with all paws placed on the ground. The maximum trial length allotted was one minute.

#### Wire Hang

Wire hang was performed to assess motor function and muscle strength in P21 male and female mice. A wire cage hopper was placed across an empty cage. Each mouse was allowed to grasp the wire cage hopper that was then carefully inverted over the empty cage. Once inverted, the latency to fall was recorded with a maximum allotted trial length of three minutes. Body weight was collected after the completion of the trial. Holding impulse was calculated as latency to fall (s) x bodyweight (kg).^43,44^

### Adult Behavioural Phenotyping

All adult neurobehavioral tests were done in male and female mice between 12-22 weeks of age (except for behavioural convulsion tracking) and conducted in the daytime when the lights were on between 7 am and 7 pm. In most cases, mice were experimentally naive at the time of behavioural testing and were not handled by experimenters prior to testing (outside of standard animal husbandry).

#### Wire Hang

Wire hang was performed in adult mice to re-assess motor function and muscle strength. A wire cage hopper was placed across an empty cage, held up by two cages on either side. Each mouse was allowed to grasp the wire cage hopper which was then subsequently inverted over the empty cage. Once inverted, the latency to fall was recorded with a maximum allotted trial length of one minute. This was immediately repeated twice more, for a total of three one-minute trials. Body weight was collected after the completion of all trials. Latency to fall was averaged across the three trials and holding impulse was calculated as average latency to fall (s) x bodyweight (kg).^43,44^

#### Open Field Test

Mice were placed individually into square chambers within a transparent plexiglass chamber (27.3 x 27.3 x 20 cm) equipped with three 16-beam IR arrays (X, Y, and Z –axis) for 2 hours. Four individual square chambers were quadrants within a larger area subdivided by a transparent plexiglass “+” insert. The entire chamber was encased in an isolated sound-attenuating chamber with dim light (30 lux). Animal activity was tracked for two hours with five-minute bins, which was analyzed by Activity Monitor Software (Med Associates, Inc.). Stereotypic activity count (given on graphs as total stereotypy number) was operationalized as the total number of beam breaks due to repetitive movements such as grooming or head bobbing.

#### Social Interaction Test

Sociability was tested using a procedure adapted from the three-chamber apparatus,^45^ in which there were no dividing walls (resulting in a singular unobstructed arena). Mice were placed in an opaque-white walled plexiglass arena (62 x 40.5 x 23 cm). The arena contained two identical inverted wire cups: one empty (non-social, NS) and one containing a novel age and sex-matched WT mouse of the same genetic background (social, S). The location of the social cup in the left vs. right side of the arena was systematically alternated between trials. Over a 10-min period, the test mouse was video recorded, and its movements were automatically tracked and analyzed using Biobserve Viewer2 software (St. Augustin, Germany). Around each inverted wire cup, a circular zone with a radius of 5 cm was defined. Sociability was measured as the time spent in the 5-cm social zone around the cup containing the novel mouse versus the time spent in the 5-cm non-social zone around the empty cup. Mice that climbed on top of either the non-social or social cup during the trial were excluded from the analysis.

#### Elevated Plus Maze

The elevated plus maze is a well-validated assay of anxiety-like behaviour in mice. In this study, the location of the mouse was tracked continuously within two “open” (25 x 5 cm) and two enclosed arms (25 x 5 x 30 cm) of an elevated arena (50 cm). The two arms of each type were arranged across from one another, extending from a central area (5 x 5 cm) in a ‘+’ configuration. Trials were conducted in dim light with light of ∼75 lux illuminating the open arms. Each mouse was placed in the central area of the elevated plus maze facing an open arm and allowed to explore the maze for 8 minutes. Each mouse was video-recorded and its movements were tracked and analyzed using Biobserve Viewer2 software (St. Augustin, Germany). The number of entries and time spent in each type of arm and the center was measured, as well as the total distance travelled in the maze. Subsequently, the percentage of time spent in the open arms was calculated by dividing the total time spent in the open arms by the total length of the test (480 seconds) and multiplying by 100.

#### Puzzle Box

The puzzle box procedure was adapted from Ben Abdallah *et al.*^46^ and was used to assess executive function and cognitive flexibility. The puzzle box arena consisted of a white opaque plexiglass box divided by a removable barrier, creating two compartments: a bright starting zone (58 cm x 28 cm) and a goal zone covered by a retractable roof (15 cm x 28 cm) that contained a mouse shelter (Lantz Enterprises Inc., Hamilton, Canada). An underpass (4 cm wide) was located beneath the barrier connecting the starting and goal zones. The experimental protocol consisted of seven trials (T1-7) over three consecutive days, with three trials per day (except for the third day). Across the trials, mice were presented with increasingly difficult obstacles constraining their path through the underpass into the goal zone. On day 1 T1, which was considered the training trial, the underpass was open and the barrier had an open door located above the underpass (4 cm x 4 cm). On T2 and T3, the doorway in the barrier was removed and the mice had to enter the goal zone exclusively via the underpass. On day 2, T4 was identical to T2 and T3, where mice must again enter the goal zone through the underpass. On T5-T6 (day 2) and T7 (day 3) the underpass was filled with clean bedding, which the mice had to dig through in order to reach the goal zone. This trial sequence allows for the testing of cognitive flexibility and problem solving through exposure to novel tasks (T2 and T5), including short- and long-term memory through repetition of specific obstacles 2 min (T3 and T6) and 24h (T4 and T7) after the initial exposure, respectively. Trials were started by placing the mouse at the end of the start zone furthest from the barrier/underpass and ended when the mouse entered the goal zone with all four paws. Upon entry to the goal box, mice were allowed to remain for 1 min. Mice were then removed and placed in a clean cage for 1 min before beginning the next trial. Mouse performance was assessed using latency to enter the goal box on each trial, with a maximum of 5 min allotted to each trial.

#### Audiogenic Seizure Testing

Audiogenic seizure testing was conducted as described in Wong *et al*..^47^ The testing apparatus consisted of a soundproof box containing a plastic mouse cage (28 x 17 x 14 cm) outfitted with a sound source (125 dB Piezo siren, electrosonic; Piezo Technologies) on the lid, which extended 5 cm into the cage. Mice aged 21-22 weeks (+/+ n = 13, males n = 8, females n = 5; Y647S/+ n = 14, males n = 4, females n = 10) were individually placed in the cage within the test apparatus and allowed to explore for a 2-min habituation period. Immediately thereafter, the sound source was played for 3 min. Video was recorded throughout the habituation and audiogenic siren sessions, and seizure activity was scored using a standard 4-point severity scale, as in Wong *et al.*.^47^

#### Behavioural Convulsion Tracking

Starting at ages 20-30 weeks, mice were monitored once weekly for 12 weeks for the presence of behavioural convulsions. During monitoring, mice were handled for one minute in a biological safety cabinet and subsequently weighed. The presence or absence of a behavioural convulsion was recorded for each mouse and if present was captured on video. If a behavioural convulsion was elicited, the mouse was allowed to recover before weighing and being returned to the home cage.

Behavioural convulsion severity was rated from videos using the categories depicted in **Supplementary Table 1**. Convulsion categories were additive, for example, the most severe convulsions included bouts of jumping and backing up in addition to characteristics seen in preceding categories.

### Pharmacological Intervention on Behavioural Convulsions

*Grin1*^Y647S/+^ mice displaying prominent convulsions and corresponding wildtype mice (age 20-30 weeks) used for 12 weeks of behavioural convulsion tracking were allocated to receive *ad libitum* access to drinking water with 6 mg/mL β-hydroxybutyrate (BHB) or drinking water alone (H_2_O control) for four weeks. An additional cohort of *Grin1*^Y647S/+^ and corresponding wildtype mice (17-26 weeks) underwent 12 weeks of behavioural convulsion tracking and were allocated to receive *ad libitum* access to 125 mg/kg lamotrigine or control chow for four weeks. After starting treatment in both cohorts, mice continued to be monitored through once-a-week handling for behavioural convulsion tracking as previously described, blinded to treatment. On the fourth week, after completing handling for behavioural convulsion tracking, both cohorts underwent testing in the open field test.

#### BHB supplementation

BHB (Real Ketones Shift, USA, 6 mg/mL) was administered orally through supplementation in water as previously described.^48^ Leak-proof bottle spouts were used on all bottles so that consumption could be monitored weekly by weighing the water bottles. Mice in the control group received water only. BHB-supplemented and control water bottles were replaced once a week.

#### Lamotrigine Diet Chow

Lamotrigine was administered orally through supplementation in chow. Lamotrigine chow (125 mg per kg of chow) was formulated in OpenStandard diet (D11112201) by Research Diets (USA). This dosage was selected based on a previous study in mice which demonstrated plasma concentrations in the human therapeutic range after subchronic treatment.^49^ Mice in the control group received the OpenStandard diet (D11112201) only.

### Statistical Analysis

Data are presented as the mean ± SEM and error bars represent SEM unless otherwise stated. Statistical analyses were performed in GraphPad Prism v. 10.1.1 (GraphPad Software, San Diego, CA) and OriginPro 9.0 (Northampton, MA, USA). Data normality was assessed, and analysis was performed using unpaired student’s t-tests (parametric) or Mann-Whitney test (non-parametric), or two-way analysis of variance (ANOVA), followed by Tukey’s multiple comparisons post-hoc test when appropriate. Power was determined using GPower (3.1.9.2). Data from male and female mice were combined for statistical analyses, except for behavioural analyses performed at P6, P21, and for analysis of adult bodyweight and holding impulse. Males and females are represented in circle and triangle symbols, respectively, when noted. For hippocampal electrophysiology, one or two slices per animal from six animals were used in each experimental group. Statistics were performed using the number of slices. Simple linear regression was used to estimate the relationship between seizure frequency and severity as a function of weeks of monitoring. Fisher’s exact test was used to analyze difference in proportions of behavioural convulsions between groups. Data analysis was not blinded.

## Results

### Clinical features of the patient carrying the Y647S/+ variant

The patient is a 34-year-old female who presented with infantile spasms and developmental delay. The patient has severe intellectual disability and requires assistance with activities of daily living. Current symptomology includes simple partial and tonic clonic seizures, low muscle tone, gastrointestinal abnormalities (constipation), sleep issues and behaviours associated with ASD. This includes repetitive interests and limited expressive language (5-10 spoken words). However, the patient demonstrates comprehension for some spoken words, instructions for simple tasks and can use body language to communicate. Caregiver reports indicate that gross motor skills are an area of patient strength, while fine motor skills, self-help and expressive communication are areas of challenge.

For seizure control, the patient’s current pharmaceutical treatment regimen (described in **Supplementary Table 2)** includes clobazam (started at age 22) and lamotrigine (started at age 26). Lamotrigine has been reported as the anti-seizure medication that best controls patient seizures after numerous trials of other medications **(Supplementary Table 3)**, although breakthrough seizures are still observed during the daytime.

### NMDARs containing GluN1-Y647S have altered sensitivity to agonists and negative modulators in oocytes

To investigate the effects of GluN1-Y647S variant subunits on NMDAR function *in vitro*, GluN1-Y647S-containing variant NMDARs or WT GluN1 were co-expressed with either WT GluN2A or GluN2B subunits in *Xenopus laevis* oocytes. Currents from oocytes expressing recombinant NMDARs were recorded under voltage clamp at -40 mV in response to the application of endogenous agonists. Concentration-response curves were generated for the NMDAR co-agonists glutamate and glycine to determine EC_50_ values and assess the potential effects of the GluN1-Y647S variant on agonist potency. For L-glutamate concentration-response studies, GluN1-Y647S-containing NMDARs displayed significantly (non-overlapping of 95% confidence intervals determined from LogEC_50_) reduced L-glutamate EC_50_ values (reflecting enhanced potency) compared to WT GluN1 when co-expressed with GluN2A (GluN1-Y647S EC_50_: 0.074 μM; WT GluN1 EC_50_: 3.2 μM) **(Fig. 1B**; **Table 1)** or GluN2B (GluN1-Y647S EC_50_: <0.01 μM; WT GluN1 EC_50_: 1.5 μM) (**Supplementary Fig. 1A; Table 1**). Similar results were found for the glycine EC_50_ of GluN1-Y647S-containing NMDARs, which was significantly reduced compared to WT GluN1 when co-expressed with GluN2A (GluN1-Y647S EC_50_: 0.036 μM; WT GluN1 EC_50_: 1.2 μM) **(Fig. 1B**; **Table 1)** and GluN2B (GluN1-Y647S EC_50_: 0.017 μM; WT GluN1 EC_50_: 0.41 μM) **(Supplementary Fig. 1B; Table 1)**. Therefore, the GluN1-Y647S variant enhances agonist potency of L-glutamate and glycine as indicated by significant reductions in EC_50_ values.

**Figure 1.**
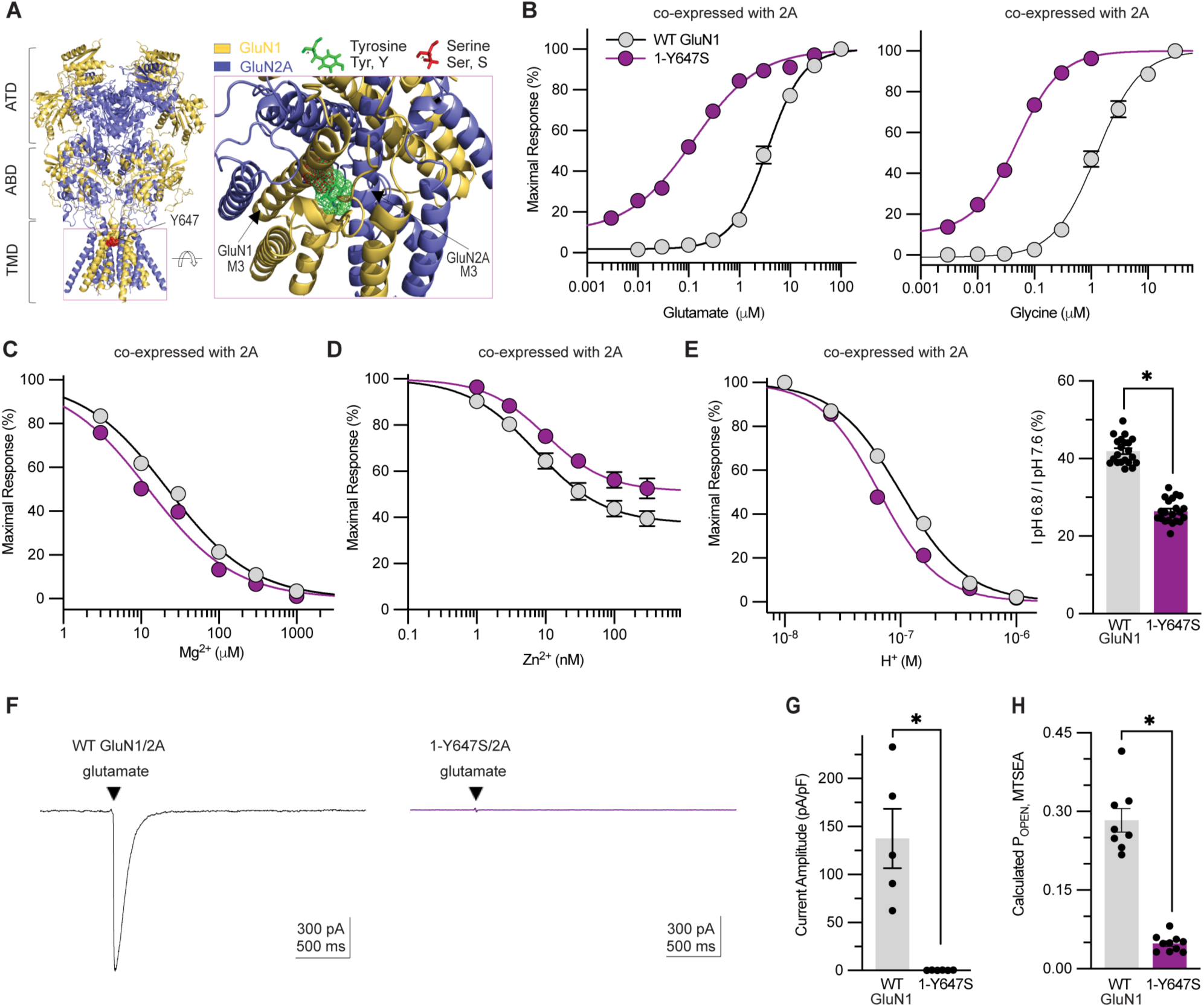
*GRIN1* p.Y647S variant alters *in vitro* NMDAR function. **(A)** The *GRIN1-*p.Y647S variant is located in M3 transmembrane domain (TMD), a critical element for channel gating. A homology model of the GluN1/GluN2A/GluN1/GluN2A receptor was built from previously published cryo-EM data for GluN1/GluN2B receptors.^93^ **(B)** Composite concentration-response curves for glutamate (*left*; in the presence of 100 μM glycine) and glycine (*right*; in the presence of 100 μM glutamate) for WT GluN1 and GluN1-Y647S co-expressed with GluN2A by TEVC oocyte recordings (V_HOLD_ -40 mV). **(C)** Composite concentration-response curves for Mg^2+^ inhibition by TEVC oocyte recordings (V_HOLD_ -60 mV). **(D)** Composite concentration-response curves for Zn^2+^ inhibition by TEVC oocyte recordings (V_HOLD_ -20 mV). **(E)** Composite concentration-response curves for proton inhibition by TEVC oocyte recordings (V_HOLD_ -40 mV). *Right*: Summary of the effects of GluN1-Y647S variant on proton sensitivity, evaluated by the ratio of current response recorded at pH 6.8 to those recorded at pH 7.6. **(F)** Representative whole cell current responses recorded under voltage clamp (V_HOLD_ -60 mV) from transiently transfected HEK cells with WT GluN1/GluN2A and GluN1-Y647S/GluN2A NMDARs are shown in response to brief 5 ms application and rapid removal of 1 mM glutamate with 100 μM glycine present in all solutions. **(G)** Summary of current amplitude. **(H)** Summary of calculated channel open probability (P_OPEN_) assessed by the degree of MTSEA potentiation by TEVC oocyte recordings (V_HOLD_ -40 mV). Data are expressed as mean ± SEM. * *p* < 0.05.

**Table 1.** Summary of the *in vitro* consequences caused by the GRIN1-p.Y647S variant.

|  | Co-expressed with GluN2A |  | Co-expressed with GluN2B |  |
| --- | --- | --- | --- | --- |
|  | WT GluN1 | GluN1-Y647S | WT GluN1 | GluN1-Y647S |
| **Glu EC<sub>50</sub> μM (n)** | 3.2 [2.6, 4.1] (11) | 0.074 [0.060, 0.090] (13) <sup>§</sup> | 1.5 [1.1, 1.9] (9) | < 0.01 (12) <sup>§</sup> |
| **Gly EC<sub>50</sub> μM (n)** | 1.2 [0.92, 1.5] (10) | 0.036 [0.028, 0.047] (12) <sup>§</sup> | 0.41 [0.30, 0.57] (6) | 0.017 [0.012, 0.025] (13) <sup>§</sup> |
| **% , pH<sub>6.8</sub> / pH<sub>7.6</sub> <sup>a</sup>** | 42 ± 0.74 (21) | 26 ± 0.66 (21) * | 15 ± 0.70 (16) | 21 ± 0.99 (8) * |
| **H<sup>+</sup> IC<sub>50</sub>, M (n)** | 1.01 E-07 (pH 7.0) (7-17) | 6.55 E-08 (pH 7.2) (7-15) | n.a. | n.a. |
| **Zinc IC<sub>50</sub>, nM (n) <sup>b</sup>** | 7.9 [5.8, 11] (8) | 9.9 [8.5, 11] (6) | n.a. | n.a. |
| **% inhibition at 300 nM zinc <sup>b</sup>** | 61 ± 3.3 (8) | 47 ± 4.4 (6) * | n.a. | n.a. |
| **Mg<sup>2+</sup> IC<sub>50</sub>, μM (n) <sup>c</sup>** | 22 [18, 27] (8) | 12 [11, 14] (6) <sup>§</sup> | 22 [16, 32] (6) | 15 [10, 22] (12) |
| **Amplitude, pA/pF** | 138 ± 31 (5) | 0.29 ± 0.08 (6) * | 49 ± 11 (5) | 0.06 ± 0.03 (6) * |
| **Calculated P<sub>OPEN</sub>** | 0.28 ± 0.02 (8) | 0.048 ± 0.005 (10)* | n.a. | n.a. |
| **Surface/total ratio (beta-lac) <sup>d</sup>** | 1.0 (5) | 0.16 ± 0.05 (5) * | 1.0 (6) | 0.44 ± 0.11 (6) * |

The concentration-response curves for glutamate were obtained in the presence of 100 μM glycine, and the glycine concentration-response curves were determined in the presence of 100 μM glutamate. The concentration-response curves for Mg<sup>2+</sup> were determined in the presence of 100 μM glutamate and glycine. Data shown are the mean EC<sub>50</sub> or IC<sub>50</sub> values with 95% confidence intervals determined from the LogEC<sub>50</sub> or LogIC<sub>50</sub> values; <sup>§</sup> indicates 95% confidence intervals that are non-overlapping with corresponding WT GluN1/GluN2A- or GluN1/GluN2B-containing NMDARs. Data are mean ± SEM for current ratio at different pH values, percentage inhibition at 300 nM zinc, current amplitude, channel open probability (P<sub>OPEN</sub>), and surface/total ratio of NMDAR expression, which were assessed by unpaired t-test (\* p < 0.05); (n) is the number of cells recorded. n.a. indicates data not available. <sup>a</sup> percentage of the current at pH 6.8 compared to that at pH 7.6; <sup>b</sup> -20 mV of holding potential; <sup>c</sup> -60 mV of holding potential; <sup>d</sup> Part of the data from Xu *et al.*<sup>26</sup> are included for comparison.

Concentration-response curves were also generated for endogenous negative allosteric modulators of the NMDAR, including extracellular Mg^2+^, Zn^2+^, and protons, to determine IC_50_ values and to assess the potential effects of the GluN1-Y647S variant on sensitivity to inhibition. The Mg^2+^ concentration-response relationship, recorded at -60 mV, revealed that the Mg^2+^ IC_50_ was significantly reduced in GluN1-Y647S-containing NMDARs compared to NMDARs that contained WT GluN1 co-expressed with GluN2A (GluN1-Y647S IC_50_: 12 μM; WT GluN1 IC_50_: 22 μM) (**Fig. 1C**; **Table 1**). The Mg^2+^ IC_50_ value for GluN1-Y647S (IC_50_ 15 μM) was comparable to WT GluN1 (IC_50_ 22 μM) when co-expressed with GluN2B (**Supplementary Fig. 1C; Table 1**). When co-expressed with GluN2A, the NMDARs containing the GluN1-Y647S variant did not display significant changes to the IC_50_ of Zn^2+^ compared to NMDARs containing WT GluN1 (GluN1-Y647S IC_50_ 9.9 nM; WT GluN1 IC_50_ 7.9 nM)**(Fig. 1D**; **Table 1**). However, the maximal degree of voltage-dependent inhibition examined in the presence of 300 nM Zn^2+^ was significantly decreased for GluN1-Y647S (% inhibition: 47 ± 4.4) compared to WT GluN1 (% inhibition: 61 ± 3.3) (**Table 1**). In addition, the influence of the Y647S variant on extracellular proton (H^+^) sensitivity was evaluated by comparing the ratio of current response recorded at pH 6.8 to pH 7.6 at a holding potential of -40 mV. When co-expressed with GluN2A, GluN1-Y647S-containing NMDARs displayed significantly enhanced inhibition at pH 6.8 (%, pH_6.8_/pH_7.6_: GluN1-Y647S 26 ± 0.66%; WT GluN1 42 ± 0.74%) (**Fig. 1E**, **Table 1**). This was consistent with the concentration-response curves for proton inhibition, which showed that Y647S variant NMDARs have an IC_50_ value corresponding to pH 7.2 (6.55×10^-8^ M), compared with pH 7.0 (1.01×10^-7^ M) for WT GluN1 co-expressed with WT GluN2A **(Fig. 1E**, **Table 1)**. However, when expressed with WT GluN2B, GluN1-Y647S had significantly reduced proton inhibition (%, pH_6.8_/pH_7.6_: GluN1-Y647S 21 ± 0.99 %; WT GluN1 15 ± 0.70 %) (**Supplementary Fig. 1D, Table 1**). Therefore, the GluN1-Y647S variant displayed increased sensitivity to Mg^2+^ channel block and decreased sensitivity to maximal inhibition by Zn^2+^. Furthermore, the GluN1-Y647S variant showed altered levels of proton inhibition, which could impact tonic levels of proton inhibition at physiological pH. Variant GluN1 showed enhanced proton inhibition when co-expressed with GluN2A, and reduced inhibition when co-expressed with GluN2B.

### NMDARs containing GluN1-Y647S affect channel biophysics and cell surface trafficking in HEK cells

To determine if the GluN1-Y647S variant alters the agonist-induced current response time course, whole cell current responses were recorded under voltage clamp at -60 mV from transiently transfected HEK cells with WT GluN1 or GluN1-Y647S co-expressed with either GluN2A or GluN2B. In response to rapid agonist application and removal to mimic synaptic events (1 mM glutamate in the presence of 100 μM glycine for 5 ms), the current amplitude of HEK cells co-expressing GluN1-Y647S was significantly reduced compared to those expressing WT GluN1 in combination with either WT GluN2A (GluN1-Y647S 0±29 ± 0.08 pA/pF; WT GluN1 138 ± 31 pA/pF) (**Fig. 1F, G**; **Table 1**) or GluN2B (GluN1-Y647S 0.06 ± 0.03 pA/pF; WT GluN1 49 ± 11pA/pF) (**Supplementary Fig. 1E, Table 1**). The amplitude of the whole cell current response mediated by NMDARs depends on several factors, including the channel open probability and receptor surface expression.^20^ Thus, these properties were evaluated to better understand the substantial (>100-fold) reduction in current response amplitude observed in HEK cells transiently transfected with GluN1-Y647S. Channel open probability of the variant GluN1 was assessed by the degree of MTSEA potentiation in TEVC oocyte recordings at a holding potential of -40 mV. When co-expressed with GluN2A, NMDARs containing the GluN1-Y647S variant displayed a significant 5-fold decrease in open probability (GluN1-Y647S calculated P_OPEN_ 0.048 ± 0.005; WT GluN1 calculated P_OPEN_ 0.28 ± 0.02) (**Fig. 1H**, **Table 1**). To evaluate whether the GluN1-Y647S variant alters receptor surface expression, a beta-lactamase (β-lac) assay was utilized to measure cell surface and total protein levels. In the assay, β-lac was fused in frame to the extracellular amino terminal domain of WT GluN1 (β-lac-GluN1-WT) or the variant GluN1-Y647S (β-lac-GluN1-Y647S). The β-lac-GluN1 fusion proteins were then co-expressed with GluN2B in HEK cells and the level of surface receptor expression was determined by β-lac cleavage of the cell-impermeable chromogenic substrate nitrocefin in the extracellular solution.^33,50^ When co-expressed with GluN2B, the GluN1-Y647S variant containing receptor displayed a 0.44 ± 0.11 surface/total ratio, relative to the 1.0 surface/total ratio assigned to WT GluN1, representing a significant reduction in NMDARs reaching the surface of the cell **(Supplementary Fig. 1F, Table 1)**. Xu *et al.*^26^ likewise found a markedly lower surface/total ratio in GluN1-Y647S variant-containing receptors compared to WT GluN1 when co-expressed with GluN2A (**Table 1, data shown for comparison**). Therefore, the GluN1-Y647S variant appears to diminish NMDAR channel open probability and surface expression, contributing to the overall reduction in whole cell current response.

### Generation and molecular characterization of *Grin1*^Y647S/+^ variant mice

To examine the consequences of the *Grin1*-Y647S (Y647S/+) variant *in vivo*, a mouse model harboring the orthologous point mutation was generated using CRISPR-Cas9 endonuclease-mediated transgenesis on a C57BL/6J genetic background **(Fig. 2A,B)**. *Grin1*^Y647S/+^ variant offspring were generated from WT x *Grin1*^Y647S/+^ breeding pairs in expected Mendelian ratios and showed normal survival, postnatal weight gain and growth from P6-10 **(Data not shown)**. Mating of *Grin1*^Y647S/+^ variant mice did not produce homozygous pups and were likely perinatal lethal based on previous studies of *Grin1* null mice^51,52^ and point mutations.^53^

**Figure 2.**
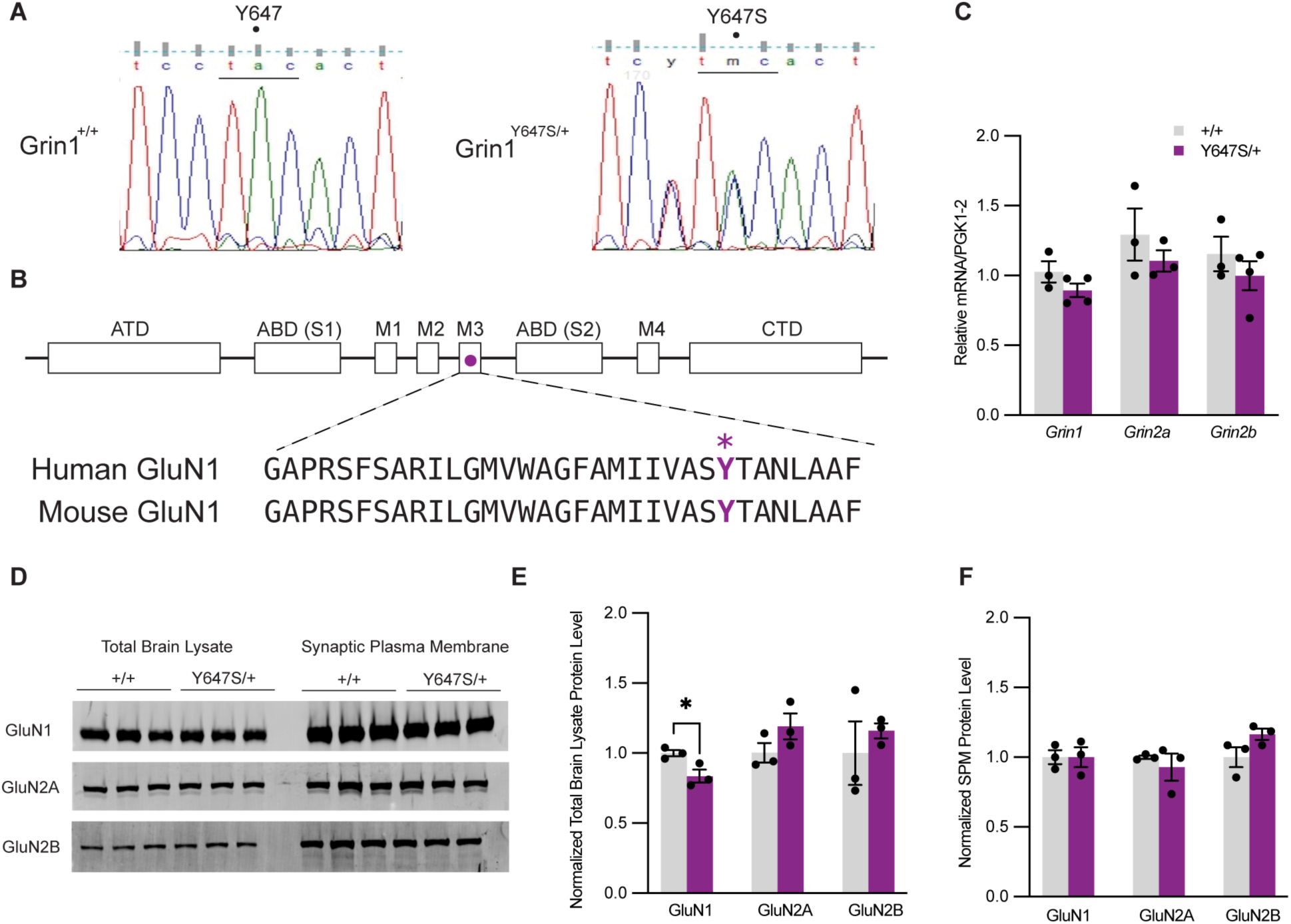
*Grin1* Y647S substitution and molecular characterization in a mouse model. **(A)** Sanger sequencing showing the A > C nucleotide mutation and diagnostic silent mutation from C > T in the preceding base pair in Y647S/+ mice. **(B)** Linear schematic representation of the GluN1 subunit, showing evolutionary conversation of amino acids between human and mouse and the site of mutation in the M3 transmembrane denoted with a circle and altered amino acid residue denoted with an asterisk. The tyrosine residue at position 647 is highly conserved across most of Vertebrata species and all four GluN2 subunits, indicating a potential critical role in channel function. ATD = amino terminal domain; CTD = carboxy terminal domain; M1-4 = transmembrane domains; agonist-binding domain (ABD) S1 and S2. **(C)** Reverse transcription PCR (RT-PCR) analysis of relative quantification of *Grin1*, *Grin2a* and *Grin2b* mRNA expression in forebrain normalized to housekeeping gene PGK1-2 (*n* =3-4) and +/+. **(D)** Western blot representation of total brain lysate and synaptic plasma membrane (SPM) fractions probed for GluN1, GluN2A, GluN2B (*n* = 3). **(E)** Relative GluN1, GluN2A, and GluN2B protein in total brain lysate normalized to total protein stain and +/+ (*n* =3; unpaired t-test: *p* = 0.0361). **(F)** Relative GluN1, GluN2A and GluN2B protein in synaptic plasma membrane fractions normalized to total protein stain and +/+ (*n* =3). Data are expressed as mean ± SEM. \**p* < 0.05.

Abundance of *Grin1* as well as *Grin2a* and *2b* mRNA in the forebrain of adult WT and *Grin1*^Y647S/+^ mice were not statistically different, indicating that the Y647S/+ mouse variant had no effect on gene transcripts for these NMDAR subunits **(Fig. 2C)**. Western blot protein analysis of whole brain from adult *Grin1*^Y647S/+^ mice indicated that GluN1 protein levels were modestly but significantly lower compared to WT mice (*p* = 0.0361), whereas GluN2A and GluN2B levels were unaffected **(Fig. 2D,E; Supplementary Fig. 2 uncropped)**. However, GluN1, 2A, and 2B protein expression in purified synaptic membrane of *Grin1*^Y647S/+^ mice were equivalent to WT **(Fig. 2D, F; Supplementary Fig. 2 uncropped)**. These results suggest that, apart from a modest reduction in whole brain GluN1 level, the heterozygous Y647S variant has minimal impact on abundance of GluN1, 2A, and 2B protein and their corresponding mRNAs.

### Deficiency in hippocampal NMDAR-mediated synaptic transmission and long-term potentiation

NMDARs contribute to cellular excitability and regulate synaptic strength; hence we used *ex vivo* hippocampal CA1 field potential recordings in hippocampal slices to identify differences in synaptic transmission and long-term potentiation for adult WT versus *Grin1*^Y647S/+^mice **(Fig. 3A)**. Fibre volley (FV) amplitude was measured in response to step wise increases in stimulus current intensity. The corresponding slopes of the I/O curves for FV vs. stimulus intensity did not significantly differ between WT and *Grin1*^Y647S/+^ mice, demonstrating largely unaffected presynaptic axon activation in *Grin1*^Y647S/+^ mice **(Fig. 3B)**. Similarly, paired-pulse facilitation, which is inversely related to neurotransmitter release probability, did not differ significantly between WT and *Grin1*^Y647S/+^ mice **(Fig. 3D)**. Together these results suggest that no major alterations in presynaptic function are caused by the *Grin1*-Y647S variant in the hippocampal CA3-CA1 Schaffer-collateral pathway.

**Figure 3.**
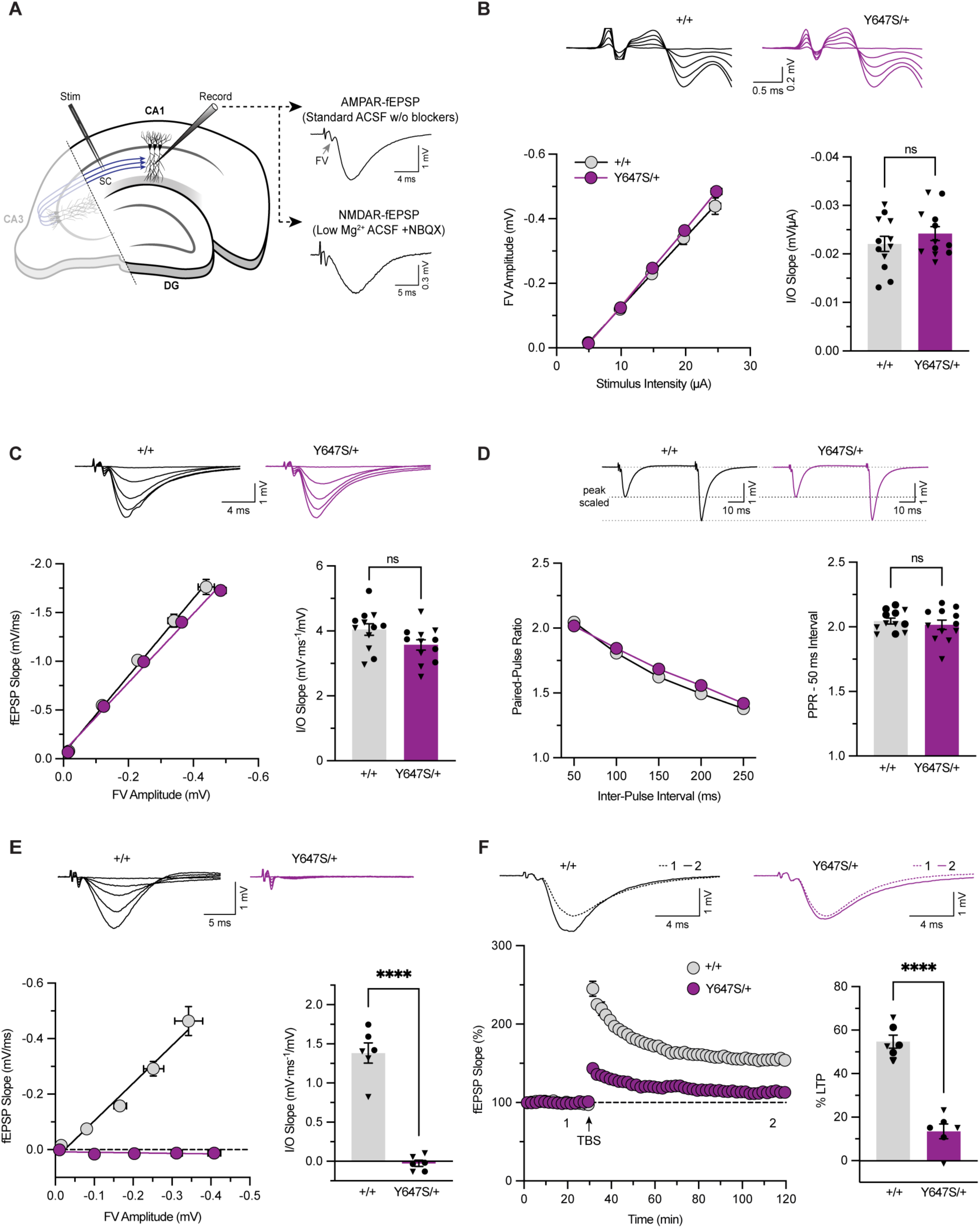
Adult *Grin1*^Y647S/+^ mice display alterations in NMDAR-mediated synaptic transmission and LTP in the hippocampus. **(A)** Schematic of a hippocampal slice showing the positions of the stimulating and recording electrodes. CA3-CA1 fibre volley (FV; indicated by a grey arrow) and AMPA receptor-mediated fEPSPs (AMPAR-fEPSP) were recorded in standard ACSF (containing 1 mM Mg^2+^) without channel blockers. NMDA receptor-mediated fEPSPs (NMDAR-fEPSP) were recorded in low (0.1 mM) Mg^2+^ ACSF with NBQX (10 µM) added to block AMPAR transmission. **(B)** Input/output (I/O) curves showing that increasing stimulus intensity corresponded to similar increases in FV amplitude in +/+ and Y647S/+ mice (left). The slope values of the linear regression fits (I/O slope) were not significantly different between genotypes, demonstrating largely unaltered presynaptic axon activation (right) (sexes combined: +/+ *n* = 12, Y647S/+ *n* =12). Representative traces shown above. **(C)** I/O curves showing the relationship between FV amplitude and AMPAR-fEPSP slope in +/+ and Y647S/+ mice (left). The I/O slopes were not significantly different between genotypes, indicating intact AMPAR-mediated synaptic transmission (right) (sexes combined: +/+ *n* = 12, Y647S/+ *n* = 12). Representative traces shown above. **(D)** Paired-pulse facilitation of AMPAR-fEPSPs measured across a range of inter-pulse intervals in +/+ and Y647S/+ mice (left). The paired-pulse ratio (PPR) at the 50 ms interval was not significantly different between genotypes, indicating no detectable differences in neurotransmitter release probability (right) (sexes combined: +/+ *n* = 12, Y647S/+ *n* = 12). Representative traces for the 50 ms interval are shown above. **(E)** I/O curves showing the relationship between FV amplitude and NMDAR-fEPSP slope in +/+ and Y647S/+ mice (left). The I/O slope was significantly reduced in Y647S/+ mice, demonstrating deficiency in NMDAR-mediated synaptic transmission (right) (sexes combined: +/+ *n* = 6, Y647S/+ *n* = 6; unpaired t-test: *p* < 0.0001). **(F)** Time-course plot showing the change in AMPAR-fEPSP slope (plotted as % of baseline) following TBS in +/+ and Y647S/+ (left). Summary of the magnitude of LTP (% change from baseline) measured 80 – 90 min post TBS (right). LTP was significantly reduced in Y647S/+ mice (sexes combined: +/+ *n* = 6, Y647S/+ *n* = 6; unpaired t-test: *p* < 0.0001). Representative traces recorded before (1) and after (2) TBS are shown above. Error bars are ± SEM. \*\*\*\**p* < 0.0001; ns: not significant. Males and females are represented in circle and triangle symbols, respectively.

Furthermore, the *Grin1-*Y647S variant had no effect on AMPAR-mediated synaptic transmission as measured by the slope of the I/O curve plotting the AMPAR-fEPSP slope as a function of FV amplitude **(Fig. 3C)**. In contrast, recordings performed in the presence of low magnesium and the AMPAR blocker NBQX revealed significantly reduced NMDAR-fEPSP slope as function of FV amplitude (*p* < 0.0001), indicating the *Grin1*-Y647S variant caused a near-complete absence of hippocampal NMDAR-mediated synaptic transmission **(Fig. 3E)**. To assess the impact of the *Grin1*-Y647S variant on synaptic plasticity, long-term potentiation at CA3-CA1 synapses was induced using a compressed TBS protocol. The magnitude of LTP (average fEPSP slope change 80-90 min post TBS) was significantly reduced in *Grin1*^Y647S/+^ mice relative to WT mice (*p* < 0.0001), indicating substantial impairment in synaptic plasticity mechanisms, consistent with diminished NMDAR-mediated charge transfer **(Fig. 3F)**.

### Adult *Grin1*^Y647S/+^ variant mice have altered hippocampal morphology

A subset of individuals with *GRIN1*-NDD display malformations of cortical development.^3^ Therefore, the potential for gross histological changes was evaluated in the neocortex of *Grin1*^Y647S/+^ mice, as well as the hippocampal subfields including the dentate gyrus. Neocortical features including thickness **(Fig. 4C)** and density of neurons **(Fig. 4D)** were found to be unaltered in adult *Grin1*^Y647S/+^ mice. In contrast, the area of the hippocampal pyramidal cell layer was found to be reduced in *Grin1*^Y647S/+^ mice compared to WT mice (*p* = 0.0290) (**Fig. 4F)**. Cell body area and the thickness of the dentate gyrus, as well as thickness of the CA1 and CA2 hippocampal subfields, were found to be unaltered (**Fig. 4F, G**).

**Figure 4.**
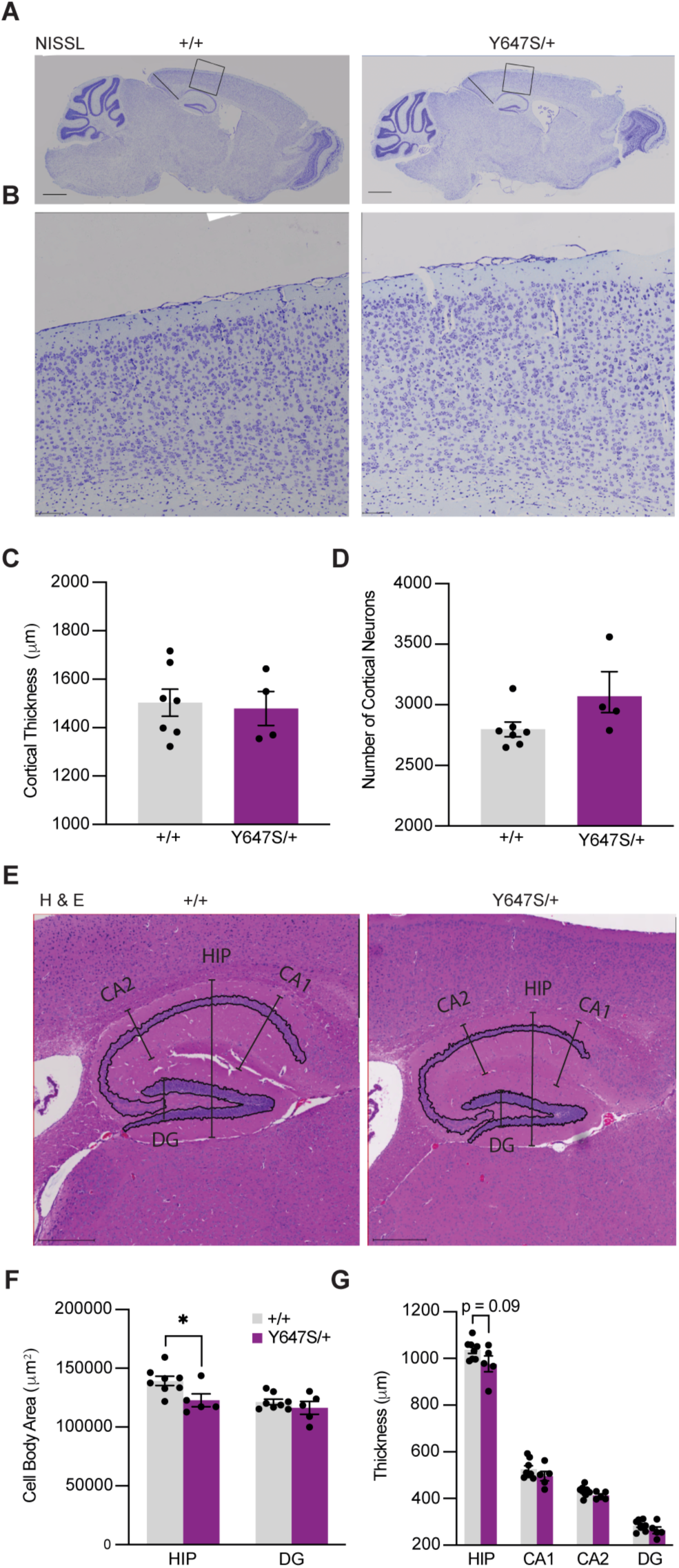
Alterations to hippocampal morphology in adult *Grin1*^Y647S/+^ mice. **(A)** Representative images of Nissl-stained brain sections with cortical thickness measurements and a 1.44 mm^2^ region used for cortical cell counting in +/+ and Y647S/+ mice (scale bars = 1 mm). **(B)** Representative images of Nissl-stained neocortex in +/+ and Y647S/+ mice (scale bars = 100 μm). (**C)** Cortical thickness and (**D)** the number of cortical neurons were not significantly different between genotypes (sexes combined: +/+ *n* = 7, Y647S/+ *n* = 4). **(E)** Representative H&E images of the hippocampus proper, CA1, CA2 and dentate gyrus (DG) thickness measurements in +/+ and Y647S/+ mice (scale bars = 400 μm). **(F)** Y647S/+ mice displayed significantly reduced area of the pyramidal cell layer in the hippocampus proper (sexes combined: +/+ *n* = 8, Y647S/+ *n* = 5; unpaired t-test: *p* = 0.0290). **(G)** Trend toward decreased thickness of the hippocampus proper (sexes combined: +/+ *n* = 8, Y647S/+ *n* = 5; unpaired t-test: *p* = 0.0874), thickness measurements averaged between both hemispheres for each mouse. Data are expressed as mean ± SEM. \**p* < 0.05.

### Alterations in communication and muscle strength in *Grin1*^Y647S/+^ postnatal development

Given the profound alterations to NMDAR function associated with the Y647S/+ variant and the severe symptomology reported in the patient including developmental delay, muscular hypotonia and limited speech capabilities, cognitive and behavioural manifestations were anticipated in the heterozygous mouse model. Thus, communicative and motor capabilities of *Grin1*^Y647S/+^ variant mice were evaluated. At P6, the righting reflex of *Grin1*^Y647S/+^ variant mice (34 ± 4.6 seconds) was present and did not significantly differ from WT (25 ± 3.6 seconds), indicating typical trunk motor control **(Fig. 5A).** However, in response to maternal isolation, P6 *Grin1*^Y647S/+^ variant pups displayed significantly more ultrasonic vocalizations in comparison to WT counterparts (two-way ANOVA main effect of genotype: F[1,64] = 8.891; *p* = 0.0041). There was a modest interaction between the effects of genotype and sex (two-way ANOVA genotype × sex interaction: F[1,64] = 3.613, *p* = 0.0618), as P6 *Grin1*^Y647S/+^ male pups appeared to display more vocalizations in comparison to WT counterparts, while female *Grin1*^Y647S/+^ did not. This may be driven by an apparent increased number of ultrasonic vocalizations in female WT compared to male WT pups **(Fig. 5B)**. At P21, *Grin1*^Y647S/+^ mice (7.6 ± 0.21 grams) displayed significantly reduced bodyweight compared to WT (8.2 ± 0.21 grams) (two-way ANOVA: main effect of genotype: F[1,60] = 4.281, *p* = 0.0428), in addition to deficits in muscle tone indicated by significantly reduced holding impulse in the wire hang task (two-way ANOVA: main effect of genotype: F[1,49] = 9.432, *p* = 0.0035) **(Fig. 5C, D**).

**Figure 5.**
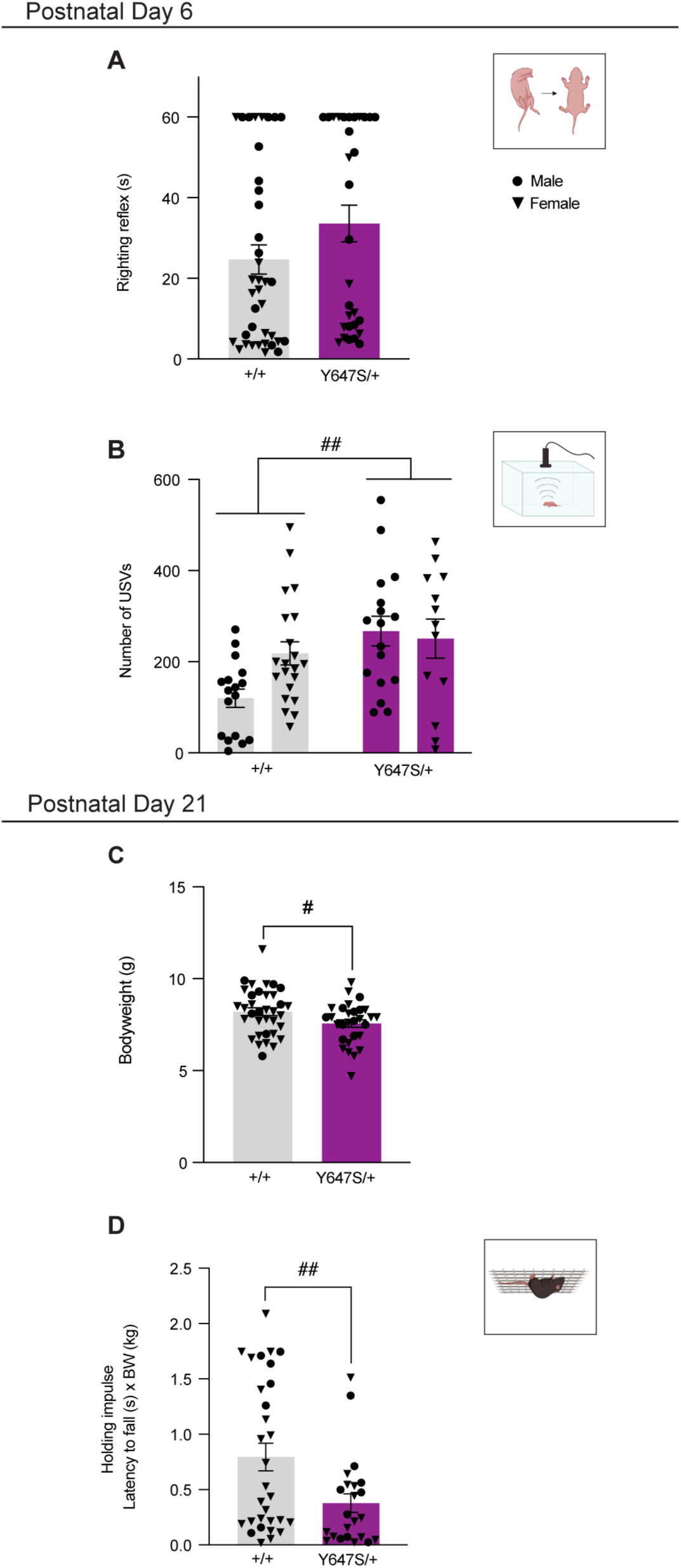
Early developmental phenotypes of *Grin1*^Y647S/+^ mice. **(A)** At P6, righting reflex was not significantly different between genotypes (+/+ *n* = 38, M = 17, F = 21; Y647S/+ *n* = 30, M = 17, F = 13) **(B)** but Y647S/+ mice displayed significantly more ultrasonic vocalizations (USVs) in response to maternal isolation compared to +/+ mice (+/+ *n* = 38, M = 17, F = 21; Y647S/+ *n* = 30, M = 17, F = 13; two-way ANOVA main effect of genotype: F[1,64] = 8.891; *p* = 0.0041). (**C)** At P21, Y647S/+ mice displayed significantly reduced bodyweight (+/+ *n* = 35, M = 10, F = 25; Y647S/+ *n* = 29, M = 10, F = 19; two-way ANOVA: main effect of genotype: F[1,60] = 4.281, *p* = 0.0428) **(D)** and Y647S/+ exhibited significantly reduced holding impulse in the wire hang task (+/+ *n* = 30, M = 7, F = 23; Y647S/+ *n* = 23, M = 9, F = 14; two-way ANOVA: main effect of genotype: F[1,49] = 9.432, *p* = 0.0035). Data are expressed as mean ± SEM. Main effect of genotype #*p* < 0.05; ##*p* < 0.01. Males and females are represented in circle and triangle symbols, respectively.

### Adult *Grin1*^Y647S/+^ variant mice display hyperactivity and recovery of muscle strength

In adulthood (12-22 weeks), there was a statistically significant interaction between the effects of genotype and sex on the bodyweight of mice (two-way ANOVA genotype × sex interaction: F[1,83] = 9.430, *p* =0.0029), with significant reductions in *Grin1*^Y647S/+^ variant bodyweight maintained in both males (WT 28 ± 0.85 grams; *Grin1*^Y647S/+^ 24 ± 0.53 grams; Tukey’s post hoc test: *p* < 0.0001) and females (WT 23 ± 0.29 grams; *Grin1*^Y647S/+^ 19 ± 0.37 grams; Tukey’s post hoc test: *p* = 0.0137) **(Fig. 6A)**. However, holding impulse in the wire hang task did not differ, indicating recovery of muscle tone deficits seen in *Grin1*^Y647S/+^ mice at P21 **(Fig. 6B)**.

**Figure 6.**
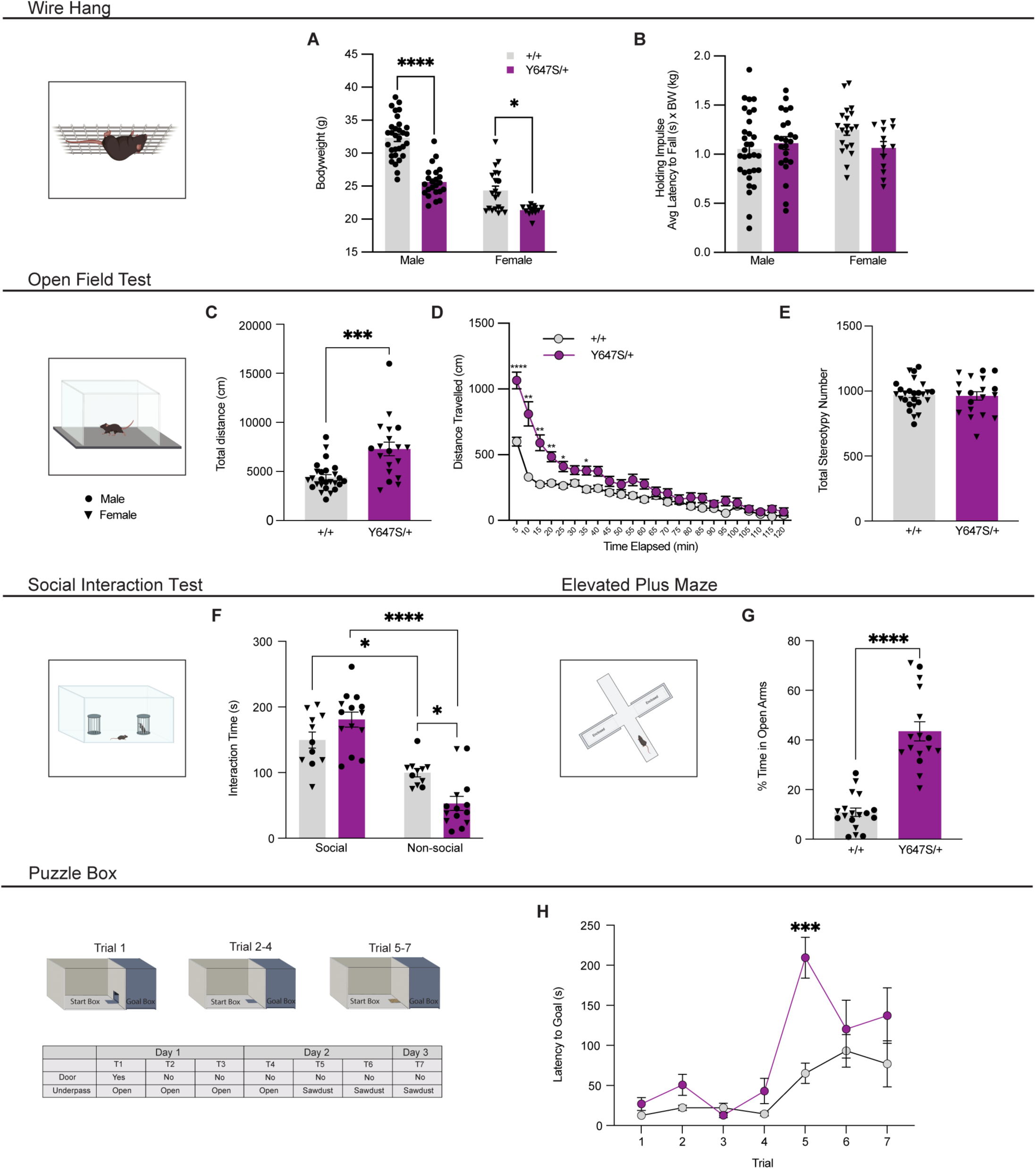
Behavioural phenotypes of adult *Grin1*^Y647S/+^ mice. **(A)** Adult (12-22 weeks) bodyweight was significantly reduced in both sexes of Y647S/+ mice compared to +/+ (+/+ *n* = 51, M = 31, F = 20; Y647S/+ *n* = 36, M = 23, F = 13; two-way ANOVA: genotype × sex interaction: F[1,83] = 9.430, *p* =0.0029; Tukey’s post hoc tests: males: *p* < 0.0001; females: *p* = 0.0137) **(B)** In adulthood, the holding impulse in the wire hang task was not significantly different between genotypes (+/+ *n* = 51, M = 31, F = 20; Y647S/+ *n* = 36, M = 23, F = 13). (**C)** Y647S/+ mice had significantly increased total distance traveled in the open field test (sexes combined: +/+ *n* = 26, Y647S/+ *n* = 19; unpaired t-test: *p* = 0.0001). **(D)** Distance traveled over time during the 2 hour open field test, with Y647S/+ mice travelling significantly greater distance in each 5-min bin from 5-25 and 35 min (sexes combined: +/+ *n* = 26, Y647S/+ *n* = 19; two-way RM-ANOVA: genotype x time interaction: F[23,989] = 16.65, *p* < 0.0001; Tukey’s post hoc tests: 5, *p* < 0.0001; 10, *p* = 0.0013; 15, *p* = 0.0014; 20, *p* = 0.0012; 25, *p* = 0.0460; 35, *p* = 0.0281). **(E)** The number of stereotypic behaviours engaged in during the open field test did not differ between genotypes (sexes combined: +/+ *n* = 26, Y647S/+ *n* = 19). **(F)** In the social interaction test, there was a statistically significant interaction between the effects of genotype and the type of stimulus (social or non-social) (sexes combined: +/+ *n* = 11, Y647S/+ *n* = 14; two-way ANOVA: genotype x stimulus interaction: F[1,46] = 13.36, *p* = 0.0007). Both genotypes spent significantly more time with the social stimulus than the non-social (Tukey’s post hoc tests: +/+ *p* = 0.0159; Y647S/+ *p* < 0.0001) and Y647S/+ spent significantly less time with the non-social stimulus compared to +/+ (*p* = 0.0169). **(G)** Percentage of time spent in the open arms of the elevated plus maze was significantly increased in the Y647S/+ compared to +/+ (sexes combined: +/+ *n* = 18, Y647S/+ *n* =16; unpaired t-test: *p* <0.0001). **(H)** There was a statistically significant interaction between the effects of genotype and trial on the latency to reach the goal box (sexes combined: +/+ *n* = 13, Y647S/+ *n* = 12; two-way ANOVA: genotype x trial interaction: F[6, 138] = 4.310, *p* = 0.0005). Y647S/+ mice displayed significantly greater latency to reach the goalbox on trial 5 where they were first presented with the puzzle box digging challenge (Tukey’s post hoc test: *p* = 0.0008). Data are expressed as mean ± SEM. \**p* < 0.05; \*\**p* < 0.01; \*\*\**p* < 0.001; \*\*\*\**p* < 0.0001. Males and females are represented in circle and triangle symbols, respectively, except in Figure 6D and 6H.

In the open field test, *Grin1*^Y647S/+^ variant mice traveled a significantly greater distance over the 2-hour trial compared to WT mice (*p* = 0.0001) **(Fig. 6C)**. There was a statistically significant interaction between the effects of genotype and time on distance traveled in the open field test (two-way ANOVA genotype × time interaction: F[23,989] = 16.65, *p* < 0.0001), with *Grin1*^Y647S/+^ variant mice traveling significantly greater distance within each of the 5-min bins between time = 5 – 25 and 35 min (Tukey’s post hoc tests: 5, *p* < 0.0001; 10, *p* = 0.0013; 15, *p* = 0.0014; 20, *p* = 0.0012; 25, *p* = 0.0460; 35, *p* = 0.0281), but not thereafter, indicating initial hyperlocomotion but intact habituation to the novel open field environment **(Fig. 6D)**. The number of stereotypic or repetitive movements exhibited during the open field test did not differ between genotypes **(Fig. 6E)**.

### Alterations in problem solving, sociability and anxiety-like behaviours in adult *Grin1*^Y647S/+^ variant mice

The social interaction test evaluates the predilection of mice to interact with a caged, same-sex stranger mouse rather than an empty cage. In this test there was a statistically significant interaction between the effects of genotype and the type of stimulus (social or non-social) (two-way ANOVA genotype × stimulus interaction: F[1,46] = 13.36, *p* = 0.0007). Both genotypes spent significantly more time with the social stimulus than the non-social (Tukey’s post hoc test: +/+ *p* = 0.0159; Y647S/+ *p* < 0.0001) **(Fig. 6F**). Interestingly, however, *Grin1*^Y647S/+^ variant mice also spent significantly less time with the non-social stimulus compared to WT (Tukey’s post hoc test: *p* = 0.0169), which may indicate exaggerated sociability **(Fig. 6F)**.

Anxiety-like behaviour can be modeled using the elevated plus maze, measured as the percentage of time spent in the open arms over the course of the session. *Grin1*^Y647S/+^ variant mice spent a significantly greater percentage of time in the open arms compared to WT mice (*p* < 0.0001), interpreted as a reduction in anxiety-like behaviour **(Fig. 6G)**.

Additionally, *Grin1*^Y647S/+^ mice were tested in the puzzle box task. In this test mice travel from a brightly lit arena to a darkened goalbox through increasingly challenging impediments to assess executive function and cognitive flexibility. There was a statistically significant interaction between the effects of genotype and trial on the latency to reach the goal box (two-way ANOVA genotype × trial interaction: F[6, 138] = 4.310, *p* = 0.0005). Comparable latencies were exhibited by *Grin1*^Y647S/+^ and WT mice during Trial 1 (T1), suggesting normal preference of each group to enter the goalbox via an open door **(Fig. 6H)**. Likewise, latencies were comparable between the genotypes during T2, the first trial in which mice must traverse an underpass to reach the goal box. However, upon first exposure to a bedding-filled underpass during T5, *Grin1*^Y647S/+^ mice were significantly slower than WT in reaching the goalbox (Tukey’s post hoc test: *p* = 0.0008) **(Fig. 6H)**. Despite atypical T5 performance, short- and long-term memory each appeared intact in variant mice, as latencies during repetition of the task 2 min (T6) and 24 h (T7) after initial exposure approached those of WT mice. Together, these results indicate that *Grin1*^Y647S/+^ mice possess typical motivation to reach the dark box and to encode and remember these trials but are initially slow to overcome novel obstacles impeding their travel.

### Adult *Grin1*^Y647S/+^ variant mice display increasingly frequent spontaneous convulsions

A prominent clinical symptom of *GRIN1*-NDD that afflicts the individual carrying the Y647S/+ variant is the presence of seizures. During weekly handling by experimenters for a duration of 12 weeks, spontaneous convulsions were completely absent in WT mice and present in *Grin1*^Y647S/+^ variant mice from the start of monitoring. Seizures persist for months in *Grin1*^Y647S/+^ variant mice with handling once each week, with 83% (19/23 mice) displaying convulsions in the 12^th^ week of monitoring (aged 32 – 42 weeks) **(Fig. 7A)**. Simple linear regression was used to test if the number of weeks of monitoring predicted the frequency of convulsions observed in the group of *Grin1*^Y647S/+^ variant mice. The fitted regression model was: number of convulsions observed in a week = (0.8322 × weeks of monitoring) + 7.591. The overall regression was statistically significant (r^2^ = 0.7502, F (1,10) = 30.03, *p* = 0.0003). Thus, the cumulative number of weekly monitoring sessions predicted the number of convulsions observed in the group of *Grin1*^Y647S/+^ mice. Convulsion severity was evaluated using the scale described in **Supplementary Table 1, Fig. 7B**, and depicted as a heat map in **Fig. 7C**. Simple linear regression was used to test if the cumulative number of weekly monitoring sessions predicted the average severity of convulsions observed in the group of *Grin1*^Y647S/+^ variant mice. The fitted regression was: average severity of convulsion = (-0.03862 × weeks of monitoring) + 3.739. The overall regression was not statistically significant (r^2^ = 0.07825, F (1,10) = 0.8490, *p* = 0.3785). Thus, the cumulative number of weekly monitoring sessions did not predict the average severity of convulsion observed in the group of *Grin1*^Y647S/+^ variant mice displaying seizures.

**Figure 7.**
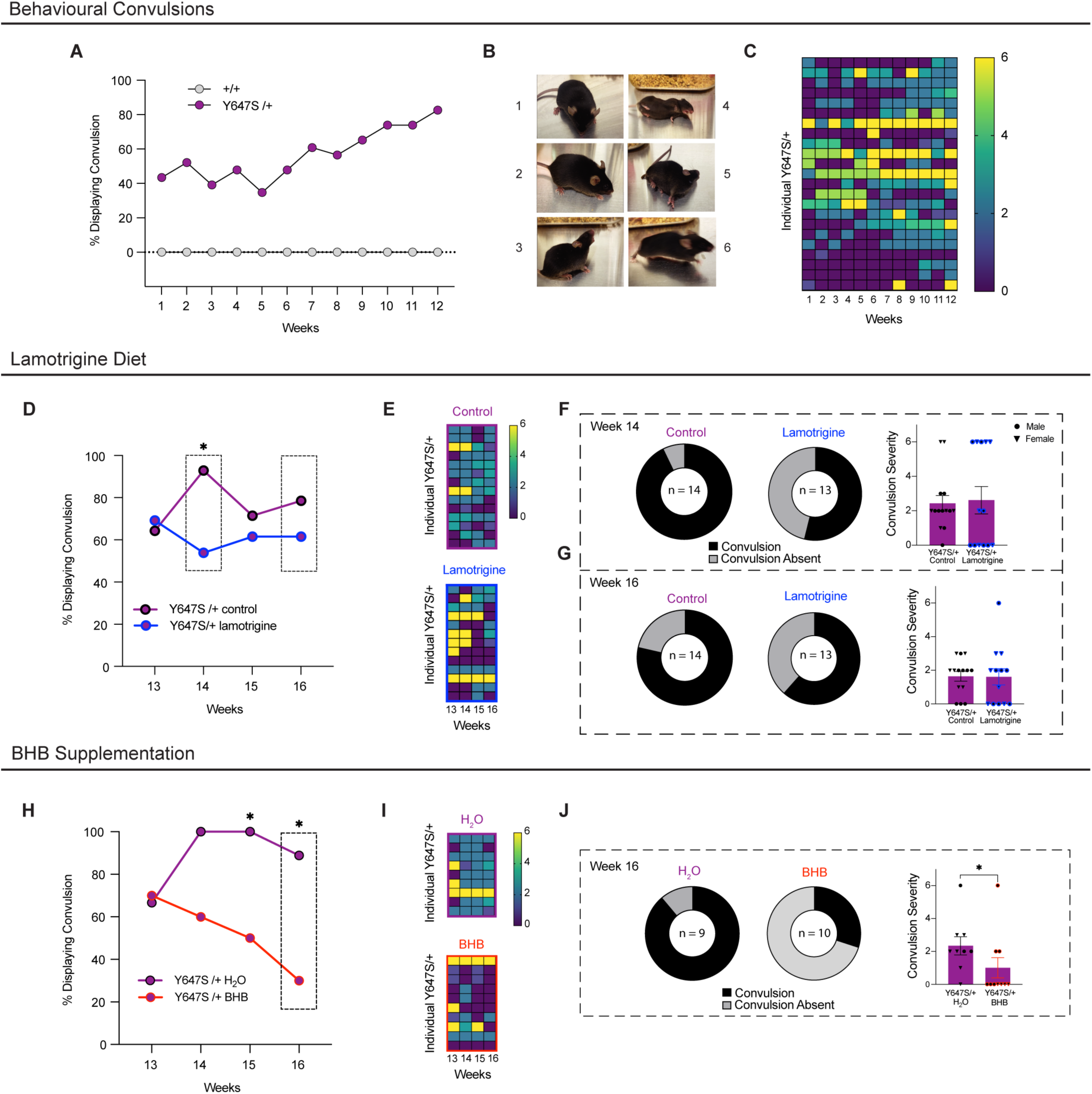
Chronic BHB Supplementation reduces behavioural convulsions in *Grin1*^Y647S/+^ mice. **(A)** Percentage of +/+ and Y647S/+ mice (age 20-30 weeks at Week 1) displaying visually identifiable convulsions over 12 weeks (sexes combined: +/+ *n* = 30, Y647S/+ *n* = 23). **(B)** Representative photographs of each category of convulsion severity observed during weekly handling described in Supplementary Table 1. **(C)** Heat map of convulsion severity observed in individual Y647S/+ mice over the course of 12 weeks (*n* = 23). **(D)** Percentage of Y647S/+ receiving 125 mg/kg lamotrigine diet or control chow displaying behavioural convulsions following initiation of treatment (Weeks 13-16). Behavioural convulsions were not observed in +/+ receiving 125 mg/kg lamotrigine or control chow and were omitted from the figure (+/+ lamotrigine *n* = 8, +/+ control *n* = 12; Y647S/+ lamotrigine *n* = 13, Y647S/+ control *n* = 14; Fisher’s exact test: Week 14: *p* = 0.0329). **(E)** Heat map of convulsion severity observed in individual Y647S/+ mice allocated to lamotrigine or control over the course of the treatment period (Weeks 13-16) (Y647S/+ lamotrigine *n* = 13, Y647S/+ control *n* = 14). **(F) Left:** Donut charts displaying the significant reduction in the proportion of Y647S exhibiting behavioural convulsions in the lamotrigine group (7/13) compared to the control group (13/14) at Week 14 (Fisher’s exact test: *p* = 0.0329). **Right:** However, lamotrigine-treated Y647S/+ mice displayed similar convulsion severity compared to those receiving control at Week 14 (Mann-Whitney test: *p* = 0.6912). **(G) Left:** Donut charts displaying the proportion of Y647S exhibiting behavioural convulsions in the lamotrigine treated group (8/13) compared to the control group (11/14) at Week 16 (Fisher’s exact test: *p* = 0.4197). **Right:** Convulsion severity of Y647S/+ in the lamotrigine-treated group compared to the control group at Week 16 (Mann-Whitney test: *p* = 0.4066). **(H)** Percentage of Y647S/+ receiving H_2_O control or BHB supplementation displaying behavioural convulsions following initiation of treatment (Weeks 13-16). Behavioural convulsions were not observed in +/+ receiving H_2_O or BHB and were omitted from the figure (+/+ BHB *n* = 9, +/+ H_2_O *n* = 14; Y647S/+ BHB *n* = 10, Y647S/+ H_2_O *n* = 9; Fisher’s exact test: Week 15: *p* = 0.0325; Week 16: *p* = 0.0198). **(I)** Heat map of convulsion severity observed in individual Y647S/+ mice allocated to H_2_O or BHB over the course of the treatment period (Weeks 13-16) (Y647S/+ BHB *n* = 10, Y647S/+ H_2_O *n* = 9). **(J) Left:** Donut charts displaying the significant reduction in the proportion of Y647S exhibiting behavioural convulsions in the BHB supplemented group (3/10) compared to the H_2_O group (8/9) at Week 16 (Fisher’s exact test: *p* = 0.0198). **Right:** BHB-supplemented Y647S/+ mice displayed significant reductions in convulsion severity compared to those receiving H_2_O at Week 16 (Mann-Whitney test: *p* = 0.0357). Convulsion severity graphs expressed as mean ± SEM. \**p* < 0.05. Males and females are represented in circle and triangle symbols, respectively in Figure 7F, G, J.

### *Grin1*^Y647S/+^ variant mice are not susceptible to audiogenic convulsions

The susceptibility of *Grin1*^Y647S/+^ variant mice to audiogenic seizure was also evaluated. When exposed to a 125 dB audiogenic sound stimulus, WT and *Grin1*^Y647S/+^ variant mice did not display susceptibility to convulsions. Only one (of a total of 14 *Grin1*^Y647S/+^ variant mice tested) displayed wild running, which was assigned the lowest severity score of 1 and hence was not categorized as a seizure.^47^

### Chronic BHB supplementation reduces the frequency of *Grin1*^Y647S/+^ spontaneous convulsions

As stated previously, lamotrigine was reported as the anti-convulsant which best controlled seizures during the daytime in the individual carrying the Y647S/+ variant. Therefore, its effectiveness in reducing behavioural convulsions was investigated in our once-a-week handling paradigm.

After 12 weeks of handling to document robust convulsions in *Grin1*^Y647S/+^ mice (**Supplementary Fig. 3A,B**), wildtype and *Grin1*^Y647S/+^ were allocated to receive 125 mg/kg lamotrigine diet chow or control diet chow. Allocation was done purposely to ensure *Grin1*^Y647S/+^ mice that displayed prominent convulsions were distributed approximately equally between lamotrigine and control treatment groups. As a result, before receiving treatment, *Grin1*^Y647S/+^ allocated to receive lamotrigine and those allocated to receive control did not differ in convulsion frequency (Fisher’s exact test: *p* = 0.3845) or severity (Mann-Whitney test: *p* = 0.9363) (**Supplementary Fig. 3C**).

After 2 weeks, *Grin1*^Y647S/+^ mice treated with lamotrigine diet chow displayed a significant reduction in the proportion of mice displaying visually identifiable behavioural convulsions compared to those receiving the control diet chow (Fisher’s exact test: *p* = 0.0329) **(Fig. 7D, E, F)**. This was not accompanied by significant changes in the severity of convulsions observed in the lamotrigine-diet treated *Grin1*^Y647S/+^ cohort compared to those receiving control diet (Mann-Whitney test: *p* = 0.6912). **(Fig. 7E, F)**. After 4 weeks, lamotrigine-treated *Grin1*^Y647S/+^ mice had a similar proportion of convulsions as those treated with the control diet (Fisher’s exact test: *p* = 0.4197) **(Fig. 7D, E, G)** and convulsion severity was similar in *Grin1*^Y647S/+^ in both groups (Mann-Whitney test: *p* = 0.4066) **(Fig. 7G)**.

At the conclusion of the study, the impact of the chronic lamotrigine diet was also explored in the open field test. Anti-seizure medicines are commonly associated with side effects of sedation, so this was an opportunity to investigate if this was observed and to examine potential effects of lamotrigine on hyperactivity observed in adult *Grin1*^Y647S/+^ mice. Distance travelled by *Grin1*^Y647S/+^ and wildtype mice was not impacted by treatment with lamotrigine (**Supplementary Fig. 5A, B**). Additionally, no treatment effects were observed in terms of the number of stereotypic behaviours engaged in during the open field test by wildtype or *Grin1*^Y647S/+^ mice **(Supplementary Fig. 5C).**

A similar study was conducted to investigate the potential anti-convulsant effects of BHB supplementation. BHB is one of the ketone bodies generated as a result of a ketogenic diet, and direct administration of BHB has been shown to exhibit anti-convulsant effects in rodent models of seizure.^54–57^

After 12 weeks of handling to document robust convulsions in *Grin1*^Y647S/+^ (**Supplementary Fig. 4A, B**), wildtype and *Grin1*^Y647S/+^ mice were allocated to receive 6 mg/mL BHB supplementation in water or water only. Allocation was done purposely to ensure *Grin1*^Y647S/+^ mice that displayed prominent convulsions were distributed approximately equally between BHB-supplemented and non-supplemented water only control groups. As a result, before receiving treatment, *Grin1*^Y647S/+^ mice allocated to receive BHB supplementation and those allocated to receive water control did not differ in convulsion frequency (Fisher’s exact test: *p* > 0.9999) or severity (Mann-Whitney test: *p* = 0.7201) **(Supplementary Fig. 4C)**.

After 3 weeks, *Grin1*^Y647S/+^ mice treated with BHB supplementation displayed a significant reduction in the proportion of mice displaying visually identifiable spontaneous convulsions compared to those receiving the water control (Fisher’s exact test: *p* = 0.0325) **(Fig. 7H, I, J)**. This reduction in convulsions remained present until the conclusion of the study after 4 weeks of treatment (Fisher’s exact test: *p* = 0.0198) **(Fig. 7H, I, J)**. At 4 weeks, the reduced convulsion frequency was accompanied by a significant reduction in the severity of convulsions observed in the BHB-supplemented *Grin1*^Y647S/+^ mice compared to those receiving water control (Mann-Whitney test: *p* = 0.0357) **(Fig. 7J)**.

At the conclusion of the study, the impact of BHB supplementation was also explored in the open field test. Distance travelled by *Grin1*^Y647S/+^ and wildtype mice was not impacted by treatment with BHB (**Supplementary Fig. 5D, E**). Additionally, no treatment effects were observed in terms of the number of stereotypic behaviours engaged in during the open field test by wildtype or *Grin1*^Y647S/+^ mice **(Supplementary Fig. 5F)**.

## Discussion

Leveraging a newly developed mouse model alongside *in vitro* methods, this study of the *Grin1* Y647S variant reveals a complex phenotype ultimately driven by diminished NMDAR function. Variant NMDARs displayed increased sensitivity to endogenous agonists of the receptor in *Xenopus* oocytes, but >300-fold decreased NMDAR response amplitudes when expressed in mammalian cell lines and no detectable synaptic NMDAR function with *ex vivo* preparations from the hippocampus of knock-in mice, despite presence of variant subunits in the synaptic plasma membrane. To understand the pathogenic nature of this variant, we examined histological and behavioural features of the *Grin1*^Y647S/+^ mice and found they displayed alterations in domains relevant to patient symptomology and to the broader characteristics of *GRIN1*-NDD. In particular, *Grin1*^Y647S/+^ mice displayed a robust spontaneous convulsion phenotype, which provided the opportunity to test one of the patient’s anti-seizure medicines (lamotrigine) and a novel treatment (BHB). Lamotrigine, the medicine that provides the best seizure management for the patient carrying the Y647S/+ variant, failed to improve spontaneous convulsion behaviour in *Grin1*^Y647S/+^ mice at the conclusion of the study. However, chronic BHB supplementation reduced the frequency and severity of spontaneous convulsions in *Grin1*^Y647S/+^ mice. Thus, BHB supplementation represents a potential anticonvulsant treatment for the *GRIN1*-NDD population and warrants continued investigation. Ultimately, the present work reveals the complex manifestations of NMDAR variation underlying *GRIN1*-NDD, identifies a potential novel treatment and provides a representative whole-organism system to further probe pathological mechanisms and test additional therapies.

### The Y647S variant imparts complex alterations to NMDAR expression and function

Our *in vitro* characterization of the Y647S variant revealed 30- to 150-fold enhanced potency to co-agonists L-glutamate and glycine and mixed effects of the variant receptor to endogenous modulators, which were in some cases dependent on the expression of specific GluN2 subunits. The Y647S variant displayed decreased sensitivity to maximal inhibition by Zn^2+^, increased sensitivity to Mg^2+^ when co-expressed with GluN2A and opposing levels of proton inhibition depending on the co-expressed GluN2 subunit. Interestingly, the complex profile of NMDAR sensitivity to agonists and endogenous modulators imparted by the Y647S variant was nearly identical to that reported for the p.Tyr647Cys (Y647C) variant.^17,16^ In both cases, structural modelling of the Y647S and Y647C variants found increased number of hydrogen bonds between the glycine ligand and the glycine-binding residues of the variant GluN1 compared to WT, offering a potential mechanism for the increased agonist affinity^17^ which was also reported in the p.Tyr647His variant (numbered as p. Tyr668His).^58^ When expressed in HEK cells, Y647S variant NMDARs had dramatically reduced (>300-fold) current amplitude in response to application and rapid removal of agonists. This is due in part to a 5-fold reduction in channel opening probability and a 2-fold reduction in cell-surface expression of Y647S variant NMDARs. This suggests that the structural changes imparted by the Y647S variant may stabilize a closed receptor state or increase the activation energy required for opening in addition to altering the trafficking of receptors to the plasma membrane, which has been noted in previous studies.^25,26^ Despite an increased *in vitro* sensitivity to agonists, the large reduction in surface expression of the Y647S variant NMDAR together with the reduced open probability will clearly reduce charge transfer mediated by variant NMDARs and thus likely accounts for the lack of observable currents.

Here, the *in vitro* trafficking deficits conferred by the Y647S variant are further elucidated in *ex vivo* hippocampal sections obtained from *Grin1^Y^*^647S/+^ mice. Strikingly, brains from variant mice show a near-complete absence of NMDAR-mediated synaptic responses in CA1 pyramidal cells and a severe deficit in NMDAR-dependent long-term potentiation measured at the Schaffer collateral-CA1 synapse. Despite this dramatic reduction of NMDAR function at this hippocampal synapse, GluN1 protein levels were found to be only modestly reduced in whole-brain homogenates from *Grin1*^Y647S/+^mice and unaltered within the synaptic membrane fraction. Therefore, loss of NMDAR functionality does not appear to be driven by reductions in GluN1 abundance at the synaptic plasma membrane. A parsimonious explanation for this apparent discrepancy is that the profound 5-fold reduction of open probability observed in heterologous systems may be even stronger in neurons, eliminating native currents despite near normal levels of protein. Alternatively, Fang *et al.*^59^ identified a mechanism by which regulated internalization of NMDARs not only reduced the number of receptors present on the cell surface, but also caused an inhibition of activity of the non-internalized NMDARs remaining on the surface mediated by increased serine phosphorylation. This offers a potential explanation for the presence of NMDARs at the synaptic plasma membrane of *Grin1*^Y647S/+^ mice that lack functionality. In these mice, NMDARs with two wildtype GluN1 subunits may properly deliver to the synaptic plasma membrane, while the NMDAR containing Y647S variant GluN1 may be removed from the membrane in response to dysfunction, thereby causing an inhibition of the activity of the NMDARs that remain on the surface membrane. Y647S variant-containing NMDARs undergoing internalization would be contained in endosomes, which are thought to be present in synaptic plasma membrane fractions.^34,60^ Future studies should examine the mechanisms underlying the preservation of relevant NMDAR protein levels at the membrane and the observed deficits in synaptic NMDAR functionality.

### *Grin1*^Y647S/+^ variant mice display early developmental and adult phenotypes reminiscent of clinical characteristics of *GRIN1*-NDD

We performed a phenotypic analysis of *Grin1*^Y647S/+^ variant mice to determine if they displayed histological and behavioural alterations in domains relevant to the symptomology seen in the patient harboring this variant and the broader clinical characteristics of *GRIN1*-NDD. In most cases, *GRIN1*-NDD is caused by a heterozygous *de novo* missense variant within the *GRIN1* gene, associated with clinical features including developmental delay, intellectual disability, alterations in speech, movement disorders, muscle tone abnormalities, epilepsy, and developmental malformation of cortex.^3,17^ Indeed, the individual carrying the Y647S variant displayed seizure onset at 5 months, developmental delay, severe intellectual disability, hypotonia and cerebral atrophy with ventriculomegaly.^2^

The most common neuroanatomical abnormality observed in *GRIN1*-NDD is a subtype of malformation of cortical development known as polymicrogyria,^17^ which is characterized by excessive cortical folds (gyri) associated with thinning or loss of cortical layers.^61,62^ Polymicrogyria-associated *GRIN1* variants tend to cluster in the S2 segment of the ligand-binding domain or the M3 transmembrane domain of GluN1.^17^ While the patient carrying the *GRIN1*^Y647S/+^ variant presents with cerebral atrophy, polymicrogyria has not been identified.^2^ However, this could be because neuroanatomical reports for this patient rely on CT scans, which cannot readily detect polymicrogyria.^17^ Indeed, a patient carrying a substitution of cysteine for the tyrosine in the same location (p. Tyr647Cys; Y647C) displayed polymicrogyria detected with MRI.^17^ Additionally, atypical hippocampal anatomy has been observed in patients with *GRIN1* variants, although more variably.^3,17^ Here, the *Grin1*^Y647S/+^ mouse was found to have normal neocortical thickness and cell counts, but abnormal hippocampal morphology. This pattern resembles the *Grin2a*^S644G/+^ variant mouse line, which displays hippocampal thinning early in development but no changes to neocortical lamination.^38^ Other mouse models with NMDAR hypofunction have also been shown to express alterations to hippocampal volume with minimal differences in neocortical structure, despite NMDARs being highly expressed in this region.^7,63,64^ Therefore, major neuroanatomical symptomology of the patient carrying the Y647S variant and those of broader *GRIN1*-NDD are recapitulated in this *Grin1*^Y647S/+^ mouse model.

Hypotonia is seen in 66% of *GRIN1*-NDD patients, including the patient carrying the Y647S variant.^2,3^ This symptom was studied by measuring righting reflex and wire hang time. The righting reflex in mice requires trunk control and is considered a standard test for the homologous labyrinthine function and body righting mechanisms observed in infants.^65,66^ In early development the righting reflex was found to be intact in *Grin1*^Y647S/+^ pups, however, later in development *Grin1*^Y647S/+^ variant mice displayed deficits in motor endurance in the wire hang task.

Changes to ultrasonic vocalizations (USVs), which are a form of vocal communication used by mice, were also observed.^42^ Mouse models of NDDs frequently display alterations in USVs,^42,67^ with the increased rate of USV emissions seen in the *Grin1*^Y647S/+^ variant mice pups likewise reported in a *GRIN2B* (*Grin2b*^C456Y/+^) patient variant mouse model^68^ and in mouse models of Rett syndrome^69^ and Fragile X syndrome.^70^

Interestingly, the motor deficits in the wire hang task that were identified in juvenile *Grin1*^Y647S/+^ mice were not observed in adult mice, indicating that developmental delay in motor endurance ameliorated with age. However, several behavioural abnormalities were evident in adult mice including hyperlocomotion, increased sociability, reduced anxiety-like behaviours, and deficits in executive function.

Movement disorders in individuals with *GRIN1-*NDD often manifest as dystonia and hyperkinetic movements that are more similar to chorea rather than the manifestations of hyperactivity present in Attention-Deficit/Hyperactivity Disorder.^3,71^ An endophenotype of hyperlocomotion is frequently observed in genetic mouse models with perturbations in NMDAR signaling.^72^ When studying a specific *GRIN2A* (*Grin2a*^S644G/+^) patient variant mouse model, Amador *et al.*^38^ noted similar hyperactivity, which was not observed in the individual carrying that variant. Rather than precisely recapitulating the human phenotype, altered locomotor activity could reflect a shared dysregulation of dopaminergic signaling caused by perturbation of NMDARs.^14,73–75^

The *Grin1*^Y647S/+^ variant mice exhibit behaviours consistent with hypersociability, which has been previously identified in several related neurodevelopmental disorder mouse models including Williams Syndrome and Angelman Syndrome.^76–79^ When considered in conjunction with the reduced-anxiety phenotype of *Grin1^Y^*^647S/+^ mice, increased sociability may be attributable to an impaired detection of danger which presents as social disinhibition.^80^ In human patients, muted danger signals could present as social behaviours that are not appropriately modulated by context or level of familiarity with others, such as approaching or touching strangers.^80–82^

The most striking endophenotype of *Grin1*^Y647S/+^ mice, which recapitulates patient symptoms, is the increasing frequency of spontaneous recurrent convulsions when monitored weekly for months. These convulsions confer an important aspect of face validity to the *Grin1*^Y647S/+^ mouse model, as the individual carrying this variant and ∼65% of individuals with *GRIN1*-NDD present with epilepsy, which in two-thirds of cases is refractory to conventional antiseizure treatment.^2,3^ Thus, the spontaneous nature of these convulsions provides an opportunity to understand etiologically relevant mechanisms and to test potential anticonvulsant treatments.^83^

### BHB supplementation is a potential novel treatment for seizures in *GRIN1*-NDD

Seizure management in individuals with *GRIN1*-NDD typically involves treatment with anticonvulsants.^3^ In the case of the patient carrying the Y647S/+ variant, lamotrigine was determined to manage seizures most effectively during daytime, after a number of anti-seizure medicines were determined to be ineffective. The presence of spontaneous convulsions in *Grin1*^Y647S/+^mice provided an opportunity to test the effectiveness of lamotrigine and assess predictive validity of this model. After 2 weeks of treatment, *Grin1*^Y647S/+^ mice that were treated with the lamotrigine diet displayed a significant reduction in convulsions relative to those on the control diet. However, this seemed to be driven by an anomalous increase in the proportion of *Grin1^Y647S/+^*displaying convulsions in the control group during this week, rather than reduced proportions of convulsions in the lamotrigine treated group. Accordingly, by the end of the study, lamotrigine-treated mice displayed convulsion frequency and severity similar to those in the control group. This was unexpected given that lamotrigine was identified as the anti-convulsant that best managed patient seizures, although breakthrough seizures were still observed in the patient. This may reflect the importance of concomitant administration of both lamotrigine and clobazam for the most effective seizure management, although, clobazam has not been shown to have significant pharmacokinetic interactions with lamotrigine.^84–86^ Most patients with drug-resistant epilepsies, frequently seen in NDDs, have treatment regimens characterized by polypharmacy,^87^ thus it may have been more relevant to assess the efficacy of the combination of anti-seizure medications taken by the patient. An alternative explanation is that the dosage of lamotrigine selected in this study was not sufficient to provide adequate protection against behavioural convulsions. This touches on a limitation of the present work, as it remains unknown if plasma concentrations of *Grin1*^Y647S/+^ mice treated with lamotrigine in this study reached the human therapeutic range as was observed in previous studies.^49^

However, in the same paradigm, chronic supplementation of BHB in water reduced the frequency and severity of spontaneous convulsions observed in *Grin1*^Y647S/+^ mice. These findings agree with previous preclinical studies which have shown the anticonvulsant efficacy of direct BHB administration in seizure-induction models employing NMDA,^56^ pilocarpine,^54,55^ or kainic acid.^57^ It has also been shown that levels of BHB correlate with the anti-convulsant efficacy of the ketogenic diet,^88^ which is reportedly used for seizure control in a number of *GRIN1*-NDD patients.^15^ While the ketogenic diet has not been studied specifically for use in *GRIN1*-NDD, it is a long-established treatment for epilepsy, that has shown effectiveness in many individuals that are refractory to anti-convulsant treatment.^89^ However, despite the diet’s efficacy, many patients exhibit low compliance or discontinuation due to its restrictive nature and lack of palatability.^89^ This may represent a significant barrier to use of the diet in some individuals with *GRIN1*-NDD, who display gastrointestinal abnormalities and feeding challenges.^3^ Thus, BHB supplementation represents a potential anticonvulsant treatment for this population, which may be a more feasible treatment option than the ketogenic diet and warrants continued investigation.

### Significance of a *GRIN1*-NDD mouse model

Individuals diagnosed with *GRIN1*-NDD display debilitating symptoms including intellectual disability, muscular hypotonia, feeding difficulties, and epilepsy.^3^ Current treatments focus on the management of symptoms; however, many patients are refractory to treatment, particularly those with epilepsy.^3^ Thus, understanding how unique variants influence NMDAR function *in vivo* is crucial to advancing our understanding of *GRIN1*-NDD and the identification of targeted treatment strategies. The creation and characterization of the *Grin1*^Y647S/+^ patient variant mouse, which recapitulates key features of the clinical characteristics of the disorder, is a crucial step toward this goal. Importantly, the present study developed a regimen of weekly handling that allows reliable measurement of convulsions in *Grin1*^Y647S/+^ mice without the use of electrical kindling or convulsant drugs, which provided an opportunity to test potential treatments in the present study. These spontaneous convulsions present in *Grin1*^Y647S/+^ mice are providing an opportunity for ongoing work to understand etiologically relevant mechanisms underlying seizures^90^ and to further test novel anti-seizure medications. Beyond this, the *Grin1*^Y647S/+^ mouse model can be used to test gene-based treatments such as antisense oligonucleotide therapies or CRISPR-based gene editing, which have shown promise in the treatment of rare neurodevelopment disorders and rely on testing in animal models that closely mimic disease conditions.^91^ Further, representative mouse models allow for the discovery of biomarkers, which if translatable to human patients provide a means of assessing disease pathology and treatment response in clinical trials.^92^

## Data availability

The data that support the findings of this study are available from the corresponding author upon reasonable request.

## Acknowledgements

We are grateful to acknowledge the work of Dr. Dawn E. Watkins-Chow and Dr. William J. Pavan from the National Human Genome Research Institute at the National Institutes of Health who created the *Grin1*^Y647S/+^ mouse model. In addition, we are grateful to Dr. Alex W.M. Hooper and Dr. David R. Hampson from the Department of Pharmaceutical Sciences at the University of Toronto who provided use of the audiogenic seizure equipment utilized in this study. Further we are thankful to Dr. Lindsey Fiddes from the Microscopy Imaging Lab at the Faculty of Medicine, University of Toronto who assisted with microscopy and to Nicholas Wilson and Danielle Stevens who assisted with behavioural experiments.

## Supplementary material

Supplementary material is available at *Brain* online.

